# Novel examples of NMD escape through alternative intronic polyadenylation

**DOI:** 10.1101/2025.08.27.672552

**Authors:** Maria Vlasenok, Antonina Kuznetsova, Dmitry A. Skvortsov, Olga A. Dontsova, Dmitri D. Pervouchine

**Author notes:** These authors contributed equally to the work (Maria Vlasenok, Antonina Kuznetsova).

## Abstract

The nonsense-mediated mRNA decay (NMD) surveillance system detects and selectively eliminates transcripts with premature stop codons. A stop codon is considered premature if it is followed by an exon-exon junction more than 50 nucleotides downstream. Pruning of the 3’-untranslated region containing such junctions through alternative polyadenylation may provide a mechanism of NMD escape. Here, we systematically examine a subclass of poison exons that carry a premature stop codon for the presence of polyadenylation sites in the downstream intron. Using data from the GTEx consortium, we observed that poison exons are more often followed by an active polyadenylation site compared with cassette exons. We also identified tissue-specific switches between NMD-targeted and NMD-escape isoforms in several human genes, including the vaccinia-related kinase *VRK3*, nuclear transcription factor *NFX1*, Notch pathway regulator *TM2D3*, and RNA helicase *DDX31*. Blocking the cleavage and polyadenylation sites in these genes using antisense oligonucleotides in human cells led to a switch from NMD-escape to NMD-target isoform, accompanied by a decrease in gene expression levels. This study reveals that NMD escape via alternative polyadenylation is a widespread, yet currently overlooked post-transcriptional mechanism of gene expression regulation.

## Introduction

Alternative splicing (AS) and alternative polyadenylation (APA) are two major steps of post-transcriptional pre-mRNA processing that substantially alter the aminoacid sequence of the gene product [1, 2, 3]. While AS operates by selecting different exon combinations, APA involves the use of different polyadenylation sites (PAS) resulting not only in alterations of the 3’-untranslated regions (3’UTRs) but also in the generation of proteins with different C-termini. The coordination between AS and APA is important in many biological processes including development [4, 5], tumorigenesis [6], and functioning of the nervous system [7].

Interestingly, both AS and APA carry out important roles beyond merely diversifying the proteome. APA significantly impacts mRNA stability, localization and translation through modulating the 3’UTR sequences, which contain *cis*-regulatory elements of microRNAs and RNA-binding proteins [8, 9]. AS is involved in the so-called unproductive splicing, a process in which a premature termination codon (PTC) is purposely introduced into the mRNA to trigger its degradation by the nonsense-mediated decay (NMD) pathway [10]. This way of regulation is particularly widespread among RNA-binding proteins [11]. Curiously, population genomics studies conclude that AS exerts even greater impact on NMD-induced changes in gene expression than on expanding protein diversity [12].

One of the current NMD models postulates that exon-exon junction complexes (EJCs) that are deposited on the mRNA during splicing are displaced after the first round of translation, while EJCs that remain bound outside of the reading frame serve as a signal that mRNA contains a PTC [13]. Yet, PTC-containing transcripts can evade degradation by NMD in a process called NMD escape [14, 15]. Several stress response pathways inhibit NMD by affecting the translation efficacy [16]. Inefficient translation termination at the PTC (readthrough), or reinitiation of translation downstream of the PTC, evicts downstream EJCs also leading to NMD escape, for instance in *ATRX* and *ATP7A* genes [17, 18]. Some RNA-binding proteins such as PABPC1, PTBP1 and hnRNPL are capable of coordinating with NMD factors to protect specific transcripts from degradation [19].

Interestingly, AS and APA can act synergistically to trigger NMD escape. If an exon containing a PTC (poison exon) is skipped, or if the sole intron downstream of a PTC is inefficiently spliced, the mRNA will no longer be subjected to NMD [14]. A similar outcome occurs when a PTC is followed by a PAS located within the downstream intron, because after cleavage and polyadenylation, no EJCs remain downstream of the PTC, causing it to be recognized as a normal stop codon. Only a few examples of NMD escape through APA are currently known, including translation of the truncated form of the *Tau* mRNA in Alzheimer’s disease (AD) patients [20], premature polyadenylation within the last post-PTC intron of the *HFE* gene [21], and activation of an intronic PAS downstream of a cryptic poison exon in the *CSF3R* gene [22].

NMD serves not only as an mRNA quality control system but also as a regulatory mechanism to maintain the abundances of physiological transcripts [23]. This prompted us to investigate whether NMD escape via APA might be a more widespread phenomenon than is currently believed. To address this question, we juxtaposed the catalogs of poison exons with the published PAS datasets and GTEx transcriptomic data to detect tissue-specific switches to the NMD isoform and simultaneous activation of an intronic PAS — a pattern that is expected for the proposed mechanism. We present statistical evidence for the fact that intronic polyadenylation happens more frequently after poison exons than after non-poison cassette exons, an observation implying that more cases of NMD escape through APA exist beyond the handful of currently known examples. An extension of this search paradigm towards cryptic poison exons and unannotated APA events further confirmed this tendency. As a proof of principle, we performed an experiment in human A549 and HeLa cells, in which antisense oligonucleotides (ASOs) targeting the intronic polyadenylation signal and cleavage site resulted in suppression of the APA isoform and upregulation of the NMD isoform in four human genes, for which NMD escape had not previously been reported.

## Methods

### Genome assembly, transcript annotation, and PAS

The GRCh38 assembly of the human genome and comprehensive CHESS transcript annotation v3.1.3 were downloaded from Genome Reference Consortium [24] and CHESS websites [25], respectively. The lists of PAS were obtained from the PolyASite 2.0 database [26] and from a study of short reads containing non-templated adenines (polyA reads) [27]. In the latter, the genomic coordinates of PAS from GTEx were lifted over to the GRCh38 assembly.

### RNA-seq data

The RNA-seq data from 9,423 samples of the GTEx project v8 were obtained from the dbGaP portal and aligned to the human genome assembly GRCh38 (hg38) using STAR aligner v2.7.8a in paired-end mode [28]. The results of NMD inactivation experiments in human 293T cells, iPS-bronchioalveolar organoid, HeLa and A549 cells followed by RNA-seq were obtained from the Gene Expression Omnibus under the accession numbers GSE126520, GSE217099, GSE125086 and GSE270310, respectively. The data were downloaded in FASTQ format and aligned to the GRCh38 (hg38) assembly using STAR aligner v2.7.8a in paired-end mode [28]. Sequencing results of transcripts bound to EJCs in HEK293 cells (RIPiT protocol [29]) together with the corresponding control RNA-seq datasets were obtained from GEO under the accession number GSE115788 and processed in the same way.

### Poison and cassette exons

Cassette exons were defined as exons that are included in a group of annotated protein-coding or NMD transcripts and skipped in another such group. Exons that share an upstream or downstream splice junctions with another non-terminal exon were not considered. Each cassette exon was characterized by the uniquely defined upstream, downstream and skipping splice junctions (*I*_1_, *I*_2_, and *E*, respectively, in Figure 1A). Annotated poison exons were defined as cassette exons that contain a stop codon 50 or more nts upstream of exon’s 3’-end (the 50-nt rule). Out of 12,528 cassette exons, 1,388 contained a stop codon, and of those 813 obeyed the 50-nt rule. To define alternative terminal exons, we selected the annotated terminal exons that reside in an intron and carry a stop codon, excluding cases when the terminal exon shared the *I*_1_ splice junction with another non-terminal exon (*n* = 3, 249).

**Figure 1:**
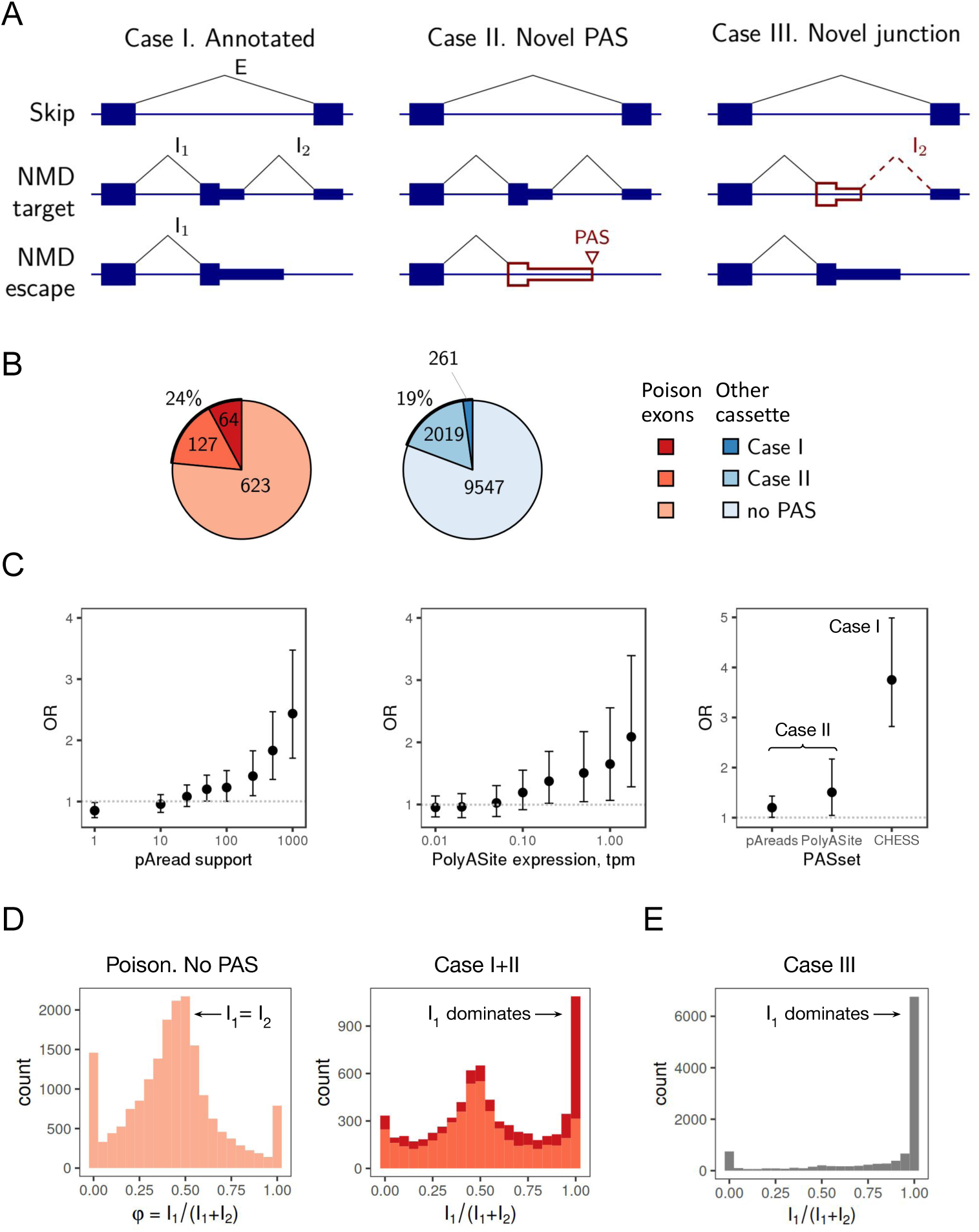
Poison exons are frequently followed by an intronic PAS. **(A)** Schematic representation of transcript isoforms in a gene containing a poison exon and a PAS within the downstream intron. In this study, we used CHESS annotations, incorporated novel PAS identified in two published databases, and included novel exon-exon junctions derived from GTEx transcriptomic data. Accordingly, a gene may contain both an annotated poison exon and PAS (Case I), or alternatively, either the PAS (Case II) or the poison exon (Case III) may be novel. Unannotated exons, PAS, and junctions are indicated with a red outline. **(B)** Number of poison and other cassette exons followed by a PAS. Color intensity indicates the annotation status of the PAS: annotated in the CHESS database (Case I), identified in the PolyASite database or predicted from poly(A) reads (Case II), or absent. The PAS derived from transcriptomic data were required to be supported by ≥50 poly(A) reads. The PAS from PolyASite were required to have an expression level of ≥0.5 TPM. **(C)** Odds ratio (OR) for the proportion of exons followed by a PAS in cassette versus poison exons. Left: *OR* as a function of polyA read support. Middle: *OR* as a function of PAS expression in TPM according to the PolyASite database. Right: *OR* for PAS identified from the PolyASite database, polyA reads, and annotated transcript ends (CHESS). Whiskers indicate 95% confidence intervals. **(D,E)** Distribution of 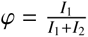 or poison exons with and without a PAS, calculated from GTEx transcriptomic data. Color coding follows the scheme in panel B.

### Read coverage and gene expression levels

The read coverage values (in BPMs, bins per million mapped reads) were calculated for each GTEx sample using the bamCoverage utility [30] considering only uniquely mapped reads. The read coverage values were averaged across samples within each detailed tissue (SMTSD) using the wiggletools utility [31]. These values were used for visualization using the Gviz package [32]. The mean read coverage was computed for 150-nt windows at the end and at the beginning of the upstream and the downstream constitutive exons flanking each cassette exon, respectively (referred to as *e*_1_ and *e*_2_). If the upstream or downstream exon was shorter than the window of 150 nts, the mean was computed over the exon’s genomic coordinates. The read counts corresponding to gene expression values were computed for each GTEX sample based on the primary GENCODE v43 annotation downloaded from the EMBL-EBI portal with featureCounts utility [33]. The read counts were normalized to TPM, and the resulting expression levels were averaged between samples within each detailed tissue type (SMTSD). In what follows, the term ‘tissue’ refers to SMTSD.

### Predicted poison exons

To predict poison exons residing inside alternative terminal exons, we extracted split reads from GTEx data using the IPSA pipeline [34]. Overall, 1,030,425 splice junctions supported by at least 10 split reads were detected, of which 332,730 were fully annotated and 402,248 had one annotated and one unannotated splice site. The latter were intersected with the list of alternative terminal exons (*n* = 3, 249) in order to select cases when (1) the acceptor splice site of the splice junction coincides with the end of the intron that contains the alternative terminal exon, and (2) the donor splice site of the splice junction is located >50 nts downstream of the annotated stop codon either within the exon or <550 nts downstream of the stop codon. The threshold of 550 nts was selected as the 95th percentile of the 3’UTR length distribution for annotated poison exons. Finally, we selected splice junctions that were collectively supported by at least 100 split reads in GTEx, which corresponds to the 10th percentile of the split read support distribution of *I*_2_ for annotated poison exons (Figure S1). This yielded 260 predicted poison exons that overlap alternative terminal exons.

### Splicing and polyadenylation metrics

To quantify exon inclusion levels, the split read counts corresponding to splice junctions (*I*_1_, *I*_2_, and *E* in Figure 1A) were obtained using the IPSA pipeline [35] and processed as described previously [34]. For the GTEx data, the split read counts were pooled within each tissue. In NMD inactivation experiments, the split read counts were pooled across biological replicates. Each exon was characterized by the following three metrics: 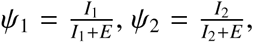 and 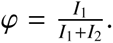 The values of these metrics with a denominator below 10 were discarded. The counts of continuous short reads overlapping the splice sites (*r*_1_, *r*_2_) were extracted by the IPSA pipeline with the default settings and analyzed using the same metrics, where *I*_1_ was replaced by *r*_1_, and *I*_2_ was replaced by *r*_2_ [35].

To quantify the level of polyadenylation at an intronic PAS, we introduced 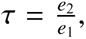 where *e*_1_ and *e*_2_ are the read coverage values in the upstream and downstream exons, respectively. This metric is sensitive to the value of the denominator, hence it was used only when *e*_1_ > 0.2, which excludes ∼40% of tissue-specific cases for all cassette exons and ∼25% for poison exons followed by a PAS (Figure S1C). By construction, *τ* can take arbitrary non-negative values, while *ψ*_1_, *ψ*_2_, and *φ* are restricted to the interval from 0 to 1. Since the values of *τ* > 1 originate from the 3’-end bias or reflect the existence of an unannotated transcript isoform, we chose to introduce an upper limit on *τ* values by redefining them as min{*τ*, 1}, that is, by setting *τ* = 1 when *τ* > 1. This cap not only allows for uniformity of the analysis but also helps to remove outliers that produce spurious tissue-specific APA events in a wrong *τ* range.

To assess splicing and polyadenylation changes, we used Δ*φ* = *φ_exp_*− *φ_ctrl_*, Δ*τ* = *τ_exp_*− *τ_ctrl_*, and Δ*ψ_i_* = *φ_i_*_,*exp*_ − *φ_i_*_,*ctrl*_, *i* = 1, 2, where ‘exp’ and ‘ctrl’ refer to EJC vs. RNA-seq experiments for RIPiT, and CHX treatment vs. untreated cells for the NMD inactivation experiments.

### Statistical analysis

The data were analyzed using R statistics software version 3.6.3. One-sided non-parametric tests were performed using normal approximation with continuity correction. For proportion comparisons, hypergeometric tests were used with Poisson or normal approximations when appropriate. The odds ratio (*OR*) values were calculated as the fraction of poison exons followed by a PAS divided by the fraction of other cassette exons followed by a PAS, across various PAS groups. Throughout the paper, *P* and *cor* denote the *P*-value and Spearman correlation coefficient, respectively. To identify significant negative correlations between *τ* and *ψ*_1_ in tissues, the respective one-sided *P*-value was estimated using asymptotic *t*-approximation. Multiple hypothesis testing was performed using Benjamini-Hochberg (BH) correction. In all figures, significance levels of 5%, 1% and 0.1% are denoted by *, ** and ***, respectively; whiskers denote 95% confidence intervals.

### Antisense oligonucleotides

ASO with 2’O methyl modified bases on a phosphorothioate backbone (2’OMe) were designed based on recommendations from [36]. Synthesis of ASO was carried out by Syntol JSC (Moscow, Russia). The sequences of used ASO are listed in Table S1.

### Cell cultures, transfections and treatments

Human A549 lung adenocarcinoma cells were maintained in Dulbecco’s modified Eagle’s medium/Nutrient Mixture F-12 (PanEco). HeLa S3 cells were cultured in Nutrient Mixture F12 medium (PanEco). Before use, the medium was supplemented with with 10% v/v fetal bovine serum (FBS), 1% GlutaMAX, 0.05 mg/ml streptomycin and 50 units/ml penicillin (all products from Thermo Fisher Scientific). Cells were cultured at 37°C in 5% CO_2_. Cells were plated at a density of 200,000 cells per well in a 12-well plate. ASO treatment was performed using Lipofectamine RNAiMAX (Invitrogen) in accordance with the manufacturer’s protocol on 70–80% confluent cells. Cells were harvested after 24h of treatment. To inactivate NMD, cycloheximide (CHX) was added to the cells 3h before harvest giving a final concentration of 300 mg/ml in the growth medium. Experiments were made in at least five independent biological replicates.

### RNA purification and cDNA synthesis

Total RNA was isolated by a guanidinium thiocyanate-phenol-chloroform method using ExtractRNA Reagent (Evrogene) in accordance with the manufacturer’s protocol. One microgram of total RNA was first subjected to RNase-free DNase I digestion (Thermo Fisher Scientific) at 37°C for 30 min to remove contaminating genomic DNA. Next, 500 ng of total RNA were used for complementary DNA (cDNA) synthesis using Magnus First Strand cDNA Synthesis Kit (Evrogene) for reverse transcription-quantitative PCR (RT-qPCR) to a final volume of 20 µl. cDNA was diluted 1:5 with nuclease-free water for quantitative PCR (qPCR) and endpoint PCR.

### Endpoint PCR and qPCR

qPCR reactions were performed in triplicates in a final volume of 12 µl in 96-well plates with 420 nM gene-specific primers and 2 µl of cDNA using 5XqPCRmix-HS SYBR reaction mix (Evrogen). Primers for qPCR are listed in Table S2. Primers for endpoint PCR are listed in Table S3. A sample without reverse transcriptase enzyme was included as a control to verify the absence of genomic DNA contamination. Amplification of the targets was carried out on CFX96 Real-Time System (Bio-Rad), with the following parameters: 95°C for 5 min, followed by 39 cycles at 95°C for 20 s, the annealing temperature (see below) for 20 s and 72°C for 20 s, ending at 72°C for 5 min. The annealing temperatures were 58°C for GAPDH and VRK3, 60°C for NFX1, DDX31 and TM2D3 (NMD-escape isoform), and 64°C for TM2D3 (NMD-target and skip isoforms). For each pair of primers in qPCR analysis, the primer efficiencies were estimated using a calibration curve. The expression of isoforms was calculated by the efficiency method with primers efficiency more than 90% in all cases.

Endpoint PCR was carried out under the following conditions: denaturing at 95°C for 5 min, 40 cycles of denaturing at 95°C for 20 s, annealing at 60°C for 20 s, and extension at 72°C for 1 min, followed by extension at 72°C for 5 min. The resulting products were analyzed on a 2% agarose gel stained with ethidium bromide and visualized using ChemiDoc XRS+ (Bio-Rad).

## Results

### PAS are more frequent downstream of poison exons

If cleavage and polyadenylation occur in the intron downstream of the PTC, the resulting transcript will lack EJCs after the stop codon. Consequently, such a PTC will be interpreted as a regular stop codon, allowing the transcript to escape NMD. We consider three scenarios for NMD-escape events driven by intronic polyadenylation. In Case I, both the poison exon and the downstream PAS are annotated (Figure 1A, left). There is a protein-coding isoform in which the poison exon is skipped (skip), and a NMD isoform that contains the poison exon (NMD target). The annotated PAS in the downstream intron introduces a third possibility, a truncated transcript, where the extended poison exon becomes an alternative terminal exon, and the PTC becomes a normal stop codon (NMD escape).

Considering that transcript catalogs are often incomplete, we also consider Case II, in which the poison exon is annotated but the intronic PAS is not (Figure 1A, middle), and Case III, in which the PAS is annotated but the poison exon is not (Figure 1A, right). To identify unannotated NMD escape isoforms in Case II, we combine evidence from PolyASite 2.0 [26] and polyA read support [27] to detect putative PAS in the intron downstream of the annotated poison exon. To identify unannotated poison exons in Case III, we rely on splice junction support derived from RNA-seq that are located more than 50 nts downstream of the stop codon and connect the annotated alternative terminal exon to the downstream constitutive exon. Such splice junctions suggest the existence of an unannotated poison exon with PTC that satisfies the 50-nt rule.

To assess the extent to which these scenarios occur, we assembled a catalog of annotated poison exons followed by an annotated PAS (Case I, *n* = 64). Poison exons were defined as cassette exons containing a stop codon that satisfies the 50-nt rule with respect to the coding frame of the major transcript isoform (a broader definition also includes generation of a PTC by a frameshift, but it is not considered here). This catalog was extended by adding the annotated poison exons followed by an intronic PAS from the PolyASite2.0 database [26] or an intronic PAS inferred from short reads with non-templated adenines (polyA reads) in GTEx [27] (Case II, *n* = 127). Finally, to detect unannotated poison exons, we considered the annotated alternative terminal exons and used split reads from GTEx to identify novel splice junctions connecting them to the downstream constitutive exon (Case III, *n* = 260). In total, this yielded 451 candidate events of NMD escape via intronic polyadenylation (Supplementary Data File 1).

As a result, 64 out 814 (7.9%) poison exons in the initial list contained an annotated PAS in the downstream intron, while only 261 out of 11,827 (2.2%) non-poison cassette exons did so (*OR* = 3.6, *P* ≈ 10^−14^) (Figure 1B, Case I). In Case II, a similar test for intronic PAS with substantial level of support (≥50 polyA reads or ≥0.5 tpm for PAS from PolyASite2.0) showed that 191 out of 814 poison exons (23.5%) were associated with a PAS in the downstream intron (Figure 1B), while only 2,280 out of 11,827 non-poison cassette cassette exons (19.3%) did so (*OR* = 1.2, *P* = 0.005). To reduce the level of noise in polyA read data, we imposed a series of stricter thresholds on the PAS support level and found even greater prevalence of PAS after poison exons, especially for PAS preceded by the canonical polyadenylation signal (Figure 1C left, Figure S2A,B). The same trend was observed when strengthening the threshold for PAS from the PolyASite2.0 database (Figure 1C, middle). Overall, metrics computed from different data sources unambiguously suggest that transcription termination occurs more frequently after poison exons than after non-poison cassette exons (Figure 1C, right).

If transcription terminates within the poison exon or in the downstream intron, the level of support by split reads for splice junctions at the 5’- and the 3’-end must be distributed differently (*I*_1_ and *I*_2_ in Figure 1A). To characterize this difference, we introduced the metric 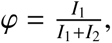, which captures the imbalance between the 5’- and the 3’-end support levels. For cassette exons, one would expect *I*_1_≈*I*_2_, and consequently, *φ* ≈ 0.5. In the extreme cases, the value *φ* ≈ 0 indicates that *I*_2_»*I*_1_ and *φ* ≈ 1 indicates that *I*_1_»*I*_2_, which is characteristic of terminal exons.

When applying this metric to human tissue transcriptomes, we found *φ* ≈ 0.5 for the majority of non-poison cassette exons (Figure S2C) and poison exons without a PAS downstream (Figure 1D, left), with a larger peak at *φ* = 0 than at *φ* = 1. A notably different pattern emerged for poison exons followed by a PAS, where the distribution displayed a pronounced peak at *φ* ≈ 1 indicating a lack of *I*_2_ reads, especially for poison exons followed by high-confidence, annotated transcript ends (Figure 1D, right). Moreover, the prevalence of *I*_1_ over *I*_2_ increased with higher PAS support (Figure S2D). Not unexpectedly, for alternative terminal exons followed by an unannotated *I*_2_ junction (Case III), we observed a stronger dominance of *I*_1_ over *I*_2_ (Figure 1E). A similar analysis using continuous reads overlapping splice sites instead of split reads revealed more frequent retention at the 3’-end of poison exons followed by a PAS, further supporting their transformation into alternative terminal exons (Figure S3).

In summary, these findings indicate that poison exons are followed by a PAS in the down-stream intron more frequently than non-poison cassette exons and that they tend to be extended at the 3’-end, thereby converting to alternative terminal exons. The combined lists for cases I, II, and III contain 451 annotated or predicted poison exons followed by an annotated or predicted PAS, representing potential NMD escape events.

### Tissue-specific switch between NMD-target and NMD-escape isoforms

To identify genes that evade NMD through intronic polyadenylation in a tissue-specific manner, we analyzed human transcriptomes from the GTEx project, focusing on tissue-specific switches between the isoform with a skipped poison exon (skip), the isoform containing the poison exon (NMD target), and the truncated isoform in which the poison exon forms part of an alternative terminal exon (NMD escape).

A widely used metric to quantify exon inclusion is the percent-spliced-in (PSI, *ψ*) ratio. It combines the split read counts from both sides of the exon 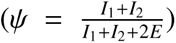 and, therefore, is not optimal for distinguishing between NMD-target and NMD-escape isoforms. Following the intron-centric approach [35], we introduced two separate metrics, 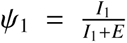 and 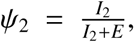, which estimate the inclusion rate at the 5’- and 3’-end of the exon separately (Figure 2A). Since the NMD-target and NMD-escape isoforms share an upstream splice junction, *ψ*_1_ measures their combined inclusion ratio, while *ψ*_2_ represents the inclusion of the poison exon alone. However, NMD-sensitive isoforms are rapidly degraded when the NMD system is active, as evidenced by the low *ψ*_2_ values in tissues (Figure S4A). Consequently, the only observable signature of NMD escape is a switch from the skip isoform to the NMD-escape isoform, both of which are stable.

**Figure 2:**
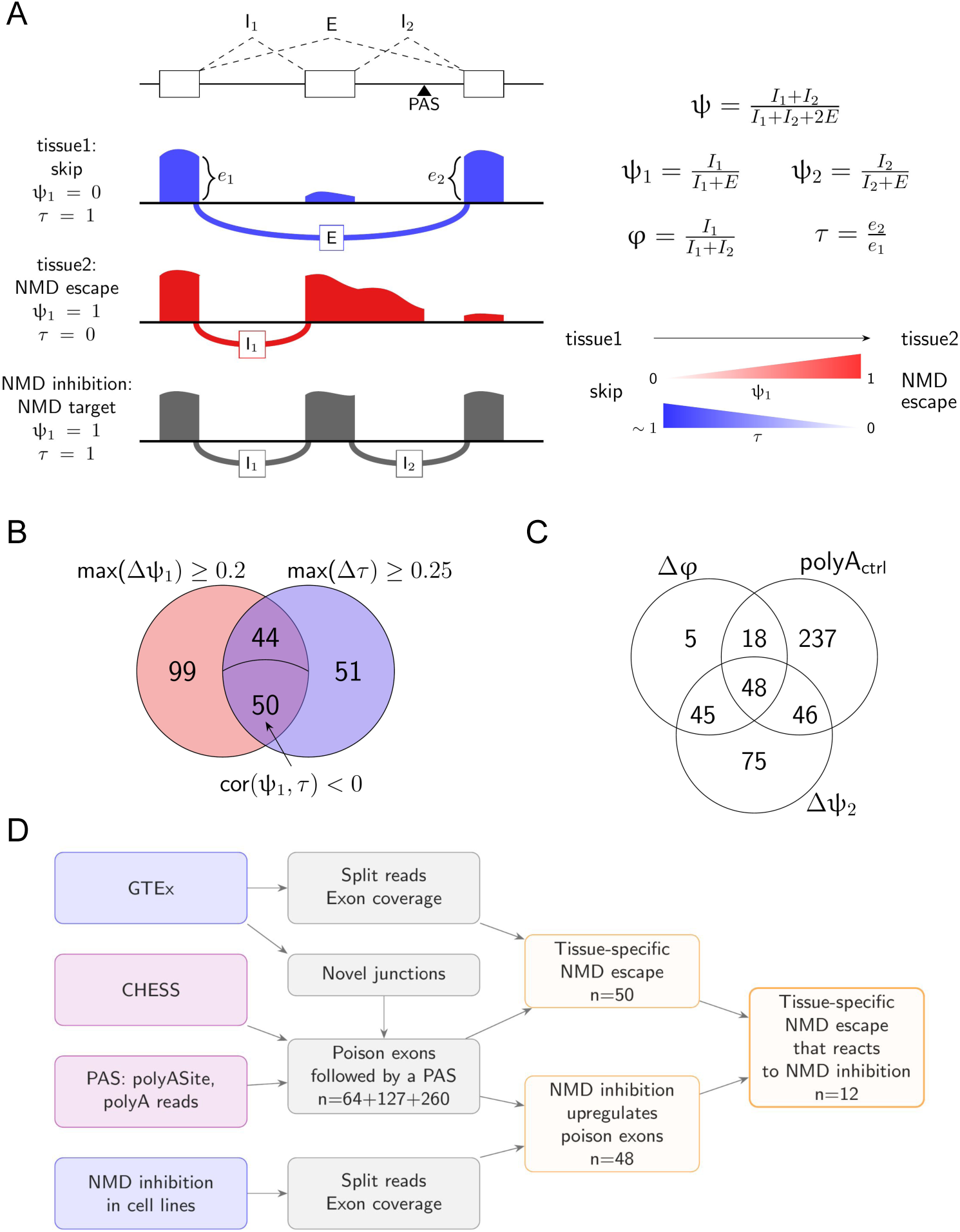
Identification of tissue-specific NMD escape events. **(A)** Left: Transcript isoforms and the metrics used for their assessment. *e*_1_ and *e*_2_ represent read coverage per nucleotide. *I*_1_, *I*_2_ and *E* denote the numbers of split reads supporting the corresponding exon-exon junctions. Right: Definition of the metrics and a schematic representation of changes in *ψ*_1_ and *τ* that are characteristic of a switch from the skip to the NMD-escape isoform: an increase in *ψ*_1_ accompanied by a decrease in *τ*. **(B)** Venn diagram of poison exons selected based on *ψ*_1_ and *τ* metrics. The portion of the intersection exhibiting a negative correlation between *ψ*_1_ and *τ* is indicated by an arrow. **(C)** Venn diagram illustrating the response of poison exons followed by a PAS to NMD inactivation. Δ indicates the changes in the respective metrics between NMD-inactivated and control conditions (see Methods). *polyA_ctrl_* refers to the category of exons with high *φ_ctrl_* or low *τ_ctrl_*values in untreated cells. **(D)** Flowchart of the analysis. Violet nodes represent database information and annotations, blue nodes correspond to transcriptomic data, gray nodes indicate processing steps, and orange nodes denote the output sets.

In addition to *ψ*_1_, we introduced another metric, *τ*, which doesn’t rely on split reads to distinguish between NMD-target and NMD-escape isoforms. It is defined as 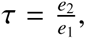, where *e*_1_ and *e*_2_ are the read coverage values in the upstream and the downstream constitutive exons, respectively (Figure 2A). This metric serves as a proxy for measuring the polyadenylation rate by the read coverage drop, which reflects the fraction of transcripts that were not cleaved and polyadenylated within the intron. A value *τ* ≈ 0 indicates a decrease in read coverage, suggesting that intronic polyadenylation predominates. Conversely, *τ* ≈ 1 indicates that the read coverage remains unchanged, implying that polyadenylation does not occur and transcription continues.

In what follows, we use the list of 451 poison exons followed by a PAS to identify cases in which an exon is skipped on one tissue but included as a terminal exon of the NMD-escape isoform in other tissues. These switches are manifested by anticorrelated tissue-specific changes in *ψ*_1_ and *τ*, where *ψ*_1_ increases, indicating splicing at the exon’s 5’-end, while *τ* decreases, indicating termination at the intronic PAS. This logic is illustrated in Figure 2B.

To select a threshold for defining tissue-specific exon inclusion, we examined the *ψ*_1_ values for cassette exons not followed by a PAS and computed their pairwise differences (Δ*ψ*_1_) between tissues (Figure S4B). The 90th percentile of this distribution was Δ*ψ*_1_ ≈ 0.2, which, when used as a threshold, identified 193 poison exons with Δ*ψ*_1_ > 0.2 in at least one pair of tissues. Similarly, to define a threshold for tissue-specific APA, we examined the *τ* values for cassette exons followed by an annotated PAS and computed their pairwise differences (Δ*τ*) between tissues (Figure S4C). A natural threshold was the 95th percentile of this distribution (Δ*τ* ≈ 0.25), which identified 145 poison exons exhibiting an absolute change in *τ* greater than 0.25 in at least one tissue pair.

The intersection of these two exon sets (max Δ*ψ*_1_ > 0.2 and max Δ*τ* > 0.25) comprised 94 exons. Imposing the additional requirement of a negative correlation between *ψ*_1_ and *τ* reduced this number to 50 exons, representing high-confidence predictions of NMD escape events (Figures 2C, S6, and S7). We retained this set for subsequent analyses because increasing Δ*τ* beyond 0.25 primarily reduces the number of candidates with minimal change in the fraction that also satisfies the other criteria, while increasing Δ*ψ*_1_ does increase this fraction, but at the expense of substantially reducing the size of the selected set (Figure S5). The characteristics of the selected exons and their associated diseases, as reported in MalaCards [37], are summarized in Table 1.

**Table 1:** Characteristics of tissue-specific NMD escape events. See also Figure S6. The table includes events in genes that are either associated with diseases reported in the literature or listed as EliteGenes for a disease in MalaCards [37]. Abbreviations: AAMR — Alacrima, achalasia, and impaired intellectual development syndrome; CNV — copy number variation.

| Gene name | range($\tau$) | range($\psi_1$) | cor($\psi_1, \tau$) | case | Associated disease |
| --- | --- | --- | --- | --- | --- |
| AKR1E2 | 0.32 | 0.26 | -0.58 | II | Cataract |
| ANKS1B | 0.67 | 0.20 | -0.81 | III | AAMR |
| ANKS1B | 0.67 | 0.96 | -0.44 | III | AAMR |
| ARMC7 | 0.31 | 0.38 | -0.58 | III | Breast Cancer (CNV) |
| BDP1 | 0.47 | 0.73 | -0.44 | II | Deafness [69] |
| BTN2A2 | 0.74 | 1.00 | -0.77 | III | Chronic Fatigue Syndrome |
| CCNH | 0.32 | 0.93 | -0.70 | III | Capillary and Arteriovenous Malformation [70, 71] |
| CDK12 | 0.33 | 0.39 | -0.64 | III | Breast, Ovarian and Prostate Cancers [72, 73, 74], Glioma [75] |
| DDX31 | 0.41 | 0.46 | -0.81 | III | Bladder Cancer [48], Pancreatic Ductal Adenocarcinoma [76] |
| FXN | 0.31 | 0.37 | -0.75 | III | Friedreich Ataxia, Atrial Standstill [77, 78], Parkinson's Disease [79] |
| GCDH | 0.33 | 0.91 | -0.42 | III | Glutaric Acidemia I [80] |
| GCOM1 | 0.41 | 0.49 | -0.69 | I | Dilated Cardiomyopathy [81] |
| LEPR | 0.31 | 0.29 | -0.63 | III | Hypogonadotropic Hypogonadism, Kallmann Syndrome [82], Leptin Receptor Deficiency and Obesity [83, 84, 85] |
| MTMR1 | 0.31 | 0.35 | -0.55 | I | Myopathy [86], Myotonic Dystrophy [87] |
| NECTIN3 | 0.70 | 0.69 | -0.92 | III | Combined Oxidative Phosphorylation Deficiency |
| NFX1 | 0.25 | 0.30 | -0.79 | III | Cervical Cancer [43, 88], Vascular Disease [89] |
| NRP1 | 0.37 | 0.48 | -0.49 | III | Covid-19 [90] |
| PDZD7 | 0.62 | 0.52 | -0.86 | III | Deafness [91], Fundus Dystrophy, Hereditary Retinal Dystrophy, Usher Syndrome [92], Nonsyndromic Hearing Loss [93] |
| PPP2R1B | 0.74 | 0.79 | -0.93 | III | Lung Cancer [94] |
| RUNDC3A | 0.34 | 0.65 | -0.72 | III | Thyroid Cancer [95], Neuroendocrine Carcinoma [96] |
| SEMA4D | 0.92 | 0.96 | -0.82 | III | Primary Sclerosing Cholangitis [97] |
| TM2D3 | 0.29 | 0.25 | -0.98 | III | Neurocardiorenal Malformation Syndrome [98] |
| TMEM45A | 0.30 | 0.24 | -0.42 | II | Glioma [99], Renal Cell Carcinoma [100], Colorectal cancer [101] |
| TPM2 | 0.58 | 0.99 | -0.89 | I | Distal Arthrogryposis [102], Cap Myopathy [103], Nemaline Myopathy [104, 105] |
| TUBGCP5 | 0.27 | 1.00 | -0.48 | II | AAMR |
| UGT1A6 | 0.39 | 0.93 | -0.65 | II | Bilirubin Metabolic Disorders including Crigler-Najjar and Gilbert Syndromes |

### Response of NMD escape events to NMD inactivation

To experimentally validate the proposed mechanism, we chose a strategy of inhibiting the production of the NMD-escape isoform using antisense oligonucleotides (ASO) that block the polyad lation signal and cleavage site. In response, we expect a switch toward the production of the NMD-target isoform and, consequently, a reduction in gene expression level. However, the isoform containing the poison exon is detectable only when the NMD system is inactivated. Therefore, we reanalyzed existing RNA-seq experiments in which NMD was inhibited using the translation inhibitor cycloheximide (CHX).

We focused on candidates expressing both NMD-escape and NMD-target isoforms containing poison exons (Figure 2C). NMD-target isoforms were selected based on Δ*ψ*_2_ (the change of *ψ*_2_ under CHX treatment), with larger Δ*ψ*_2_ indicating higher expression of the NMD-target isoform. Additionally, a decrease in the *φ* metric (Δ*φ*) under the CHX treatment was required to capture an increase in the relative abundance of NMD-target isoforms among the combined pool of NMD-target and NMD-escape isoforms. High *φ* values or low *τ* values in control experiment without CHX were used as indicators that NMD-escape isoforms are sufficiently abundant when the NMD system is active (polyA_ctrl_ in Figure 2C).

To select thresholds, we examined the set of cassette exons containing a stop codon followed by a PAS, which included both poison exons and non-poison exons with stop codons that do not satisfy the 50-nt rule. The *φ* distribution showed a pronounced peak at *φ* = 1, accompanied by a notable increase at *φ* = 0.5 upon CHX treatment for poison exons (Figures S8A). The Δ*ψ*_2_ distribution was skewed towards positive values, indicating that the inclusion rate of NMD-target isoform also increases (Figure S8B). These observations confirm that the relative abundance of the NMD-target isoform, with respect to both NMD-escape and skip isoforms, increases under NMD inhibition.

However, CHX treatment inhibits NMD by blocking the translation elongation phase, thereby affecting other pathways. An alternative approach to detect NMD-sensitive transcripts is to pull down transcripts bound to EJCs using the RIPiT protocol, which isolates post-splicing, pretranslational mRNA-EJC particles [29]. To increase the power of detecting NMD-target transcripts, we applied the same approach to analyze the RIPiT dataset obtained from HEK293 cells [29]. Although the RIPiT dataset yielded fewer genes expressing both NMD-escape and NMD-target isoforms, the results showed substantial overlap between experimental approaches (Figure S8C). For example, 56 of 85 poison exons with higher inclusion in the EJC-bound fraction also displayed increased inclusion upon CHX treatment (Δ*_CHX_ψ*_2_ and Δ*_RIPiT_ ψ*_2_ sets, *P* < 10^−14^). These findings support the robustness of our observations across different experimental methods (Figure S8D).

By combining the lists obtained for the CHX and RIPiT protocols with the results of the tissue-specific analysis, we obtained 11 genes expressing both the NMD-escape and NMD-target isoforms in cell lines and exhibiting a tissue-specific switch between them (*NFX1*, *TM2D3*, *VRK3*, *TPM2*, *BDP1*, *PITPNM2*, *DDX31*, *NMT2*, *UGT1A6*, *THAP6*, and *CDK12*). We subsequently focused on the *VRK3*, *NFX1*, *TM2D3*, and *DDX31* genes (extended correlation plots in Figure S9) as they were expressed at sufficiently high levels in the available cell lines (A549 and HeLa).

### Polyadenylation inhibition promotes NMD isoform expression

*VRK3* encodes vaccinia-related kinase 3, whose kinase activity plays a role in cell cycle regulation [38]. Additionally, *VRK3* plays a role in preventing neuronal apoptosis caused by prolonged oxidative stress, with CDK5 activating neuroprotective signaling via VRK3 phosphorylation [39].

Our analysis indicates that approximately half of the *VRK3* transcripts undergo NMD escape in cervix and esophagus, as evidenced by lower read coverage in the downstream constitutive exon, a read coverage drop at the PAS, and low *I*_2_ values (Figure 3A). In contrast, in adrenal glands and kidneys, the exon-skipping isoform predominates (*ψ*_1_ = 0.22 and *ψ*_1_ = 0.25, respectively), with no drop in read coverage in the downstream exon. This model potentially explains the increased VRK3 production in kidneys [40]. We found that *VRK3* also harbors an upstream PAS, which is preferentially activated in the brain and could represent a distinct mode of NMD escape in neural tissues (Figure S10). This observation is consistent with VRK3 supporting neuronal survival under oxidative stress via CDK5-mediated phosphorylation [39].

**Figure 3:**
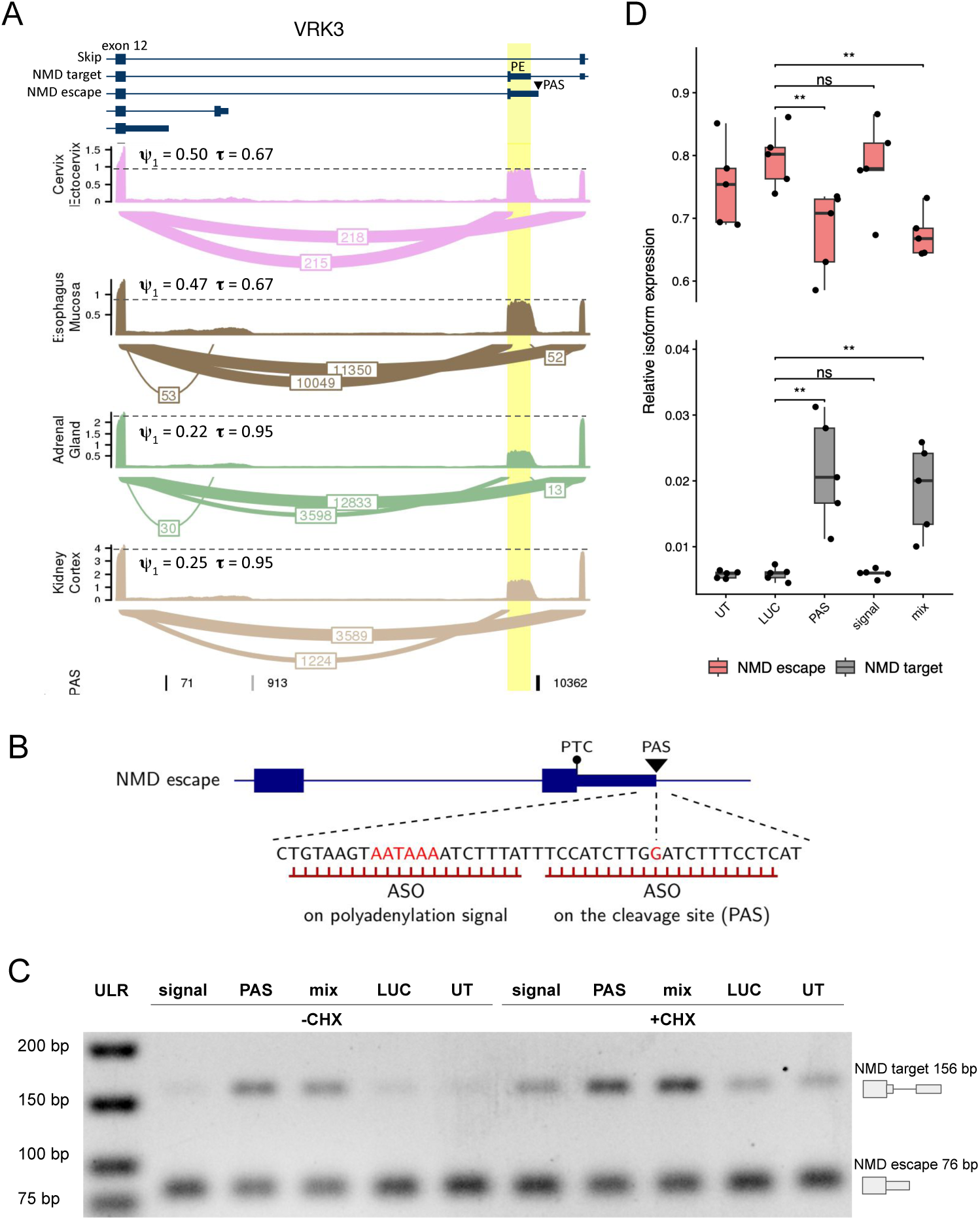
NMD escape in the *VRK3* gene. **(A)** Read coverage profiles and sashimi plots illustrating a tissue-specific switch from the skip to the NMD-escape isoform in the *VRK3* gene. The poison exon followed by a PAS is highlighted. The scale of the vertical axes represents read coverage density. Numbers on the sashimi plots indicate split read counts combined across samples from each tissue. Transcript models from CHESS are shown at the top. The PAS track indicates the number of poly(A) reads supporting each PAS. Dotted lines highlight the drop in read coverage between the upstream and downstream exons. **(B)** Genomic sequence surrounding the polyadenylation signal and the cleavage site (polyadenylation site, PAS) targeted by ASOs. **(C)** Endpoint RT-PCR analysis of untreated cells (UT), cells treated with ASO targeting luciferase (LUC), ASO targeting the polyadenylation signal (signal), ASO targeting the cleavage site (PAS), and their combination (mix), with (+CHX) or without (-CHX) cycloheximide treatment. **(D)** RT-qPCR analysis showing the fraction of NMD-escape (red) and NMD-target (grey) isoforms among all three isoforms under ASO treatment in the presence of CHX. ULR — ultra low range DNA ladder.

To test the proposed mechanism in *VRK3*, we designed ASO sequences that block the polyaden lation signal and the cleavage site through complementary base pairing (Figure 3B). These ASOs, along with a mock ASO targeting an off-target site, were transfected into human adenocarcinoma A549 cells, with and without NMD inhibition by CHX. Endpoint RT-PCR showed that transfection with 25 nM of the ASO targeting the cleavage and polyadenylation site, either alone or in combination with a second ASO blocking the polyadenylation signal, efficiently promoted inclusion of the poison exon and suppressed production of the NMD-escape isoform. In contrast, transfection with the control ASO targeting an off-target site didn’t induce the NMD-target isoform (Figure 3C). The observed effects were more significant in cells treated with CHX.

Quantitative PCR (RT-qPCR) using isoform-specific primers confirmed an upregulation of the NMD-target isoform and downregulation of the NMD-escape isoform following ASO treatment (Figure 3D). The treatment with ASO masking the PAS was more effective than with ASO targeting the polyadenylation signal. Measurement of *VRK3* mRNA levels revealed a reduction of the overall gene expression upon treatment with ASO mix in cells not exposed to CHX (Figure S11). This outcome is expected, as mRNA degradation occurs more efficiently when the NMD system is active, confirming that the fraction of *VRK3* transcripts degraded via the NMD pathway can be increased by inhibiting an intronic PAS.

The next target, *NFX1*, encodes a transcriptional repressor that binds the X-box motif in the promoter regions of major histocompatibility complex class II genes and the telomerase reverse transcriptase (hTERT) gene. It is expressed in multiple transcript isoforms with distinct functions in human epithelial cells. The truncated isoform results in NFX1-91 protein that represses hTERT transcription in keratinocytes, playing a role in epidermis function [41]. The long isoform NFX1-123, in contrast, resides in the cytoplasm and is able to increase hTERT levels post-transcriptionally via binding to 5’UTR of its mRNA [42]. NFX1-123 contributes to cellular immortalization and, unlike NFX1-91, is upregulated in cervical cancers [43]. Evidence suggests that *NFX1* may also be involved in cerebellar development regulation [44].

Our analysis identified a 279-nt cryptic poison exon located within the alternative terminal exon of *NFX1* (Figure 4A). In accordance with [44], expression of this exon was observed in the cerebellum, cerebellar hemisphere, and skeletal muscle, possibly due to lower NMD efficiency in these tissues [45]. This exon harbors two tandemly arranged acceptor splice sites separated by three nucleotides, but we treated them as a single site for simplicity. Our analysis suggests a tissue-specific switch from tissues such as the pancreas, where the NMD-target isoform is predominantly expressed, to tissues such as the skin, where both APA and overall gene expression increase, indicating NMD escape (Figure 4A).

**Figure 4:**
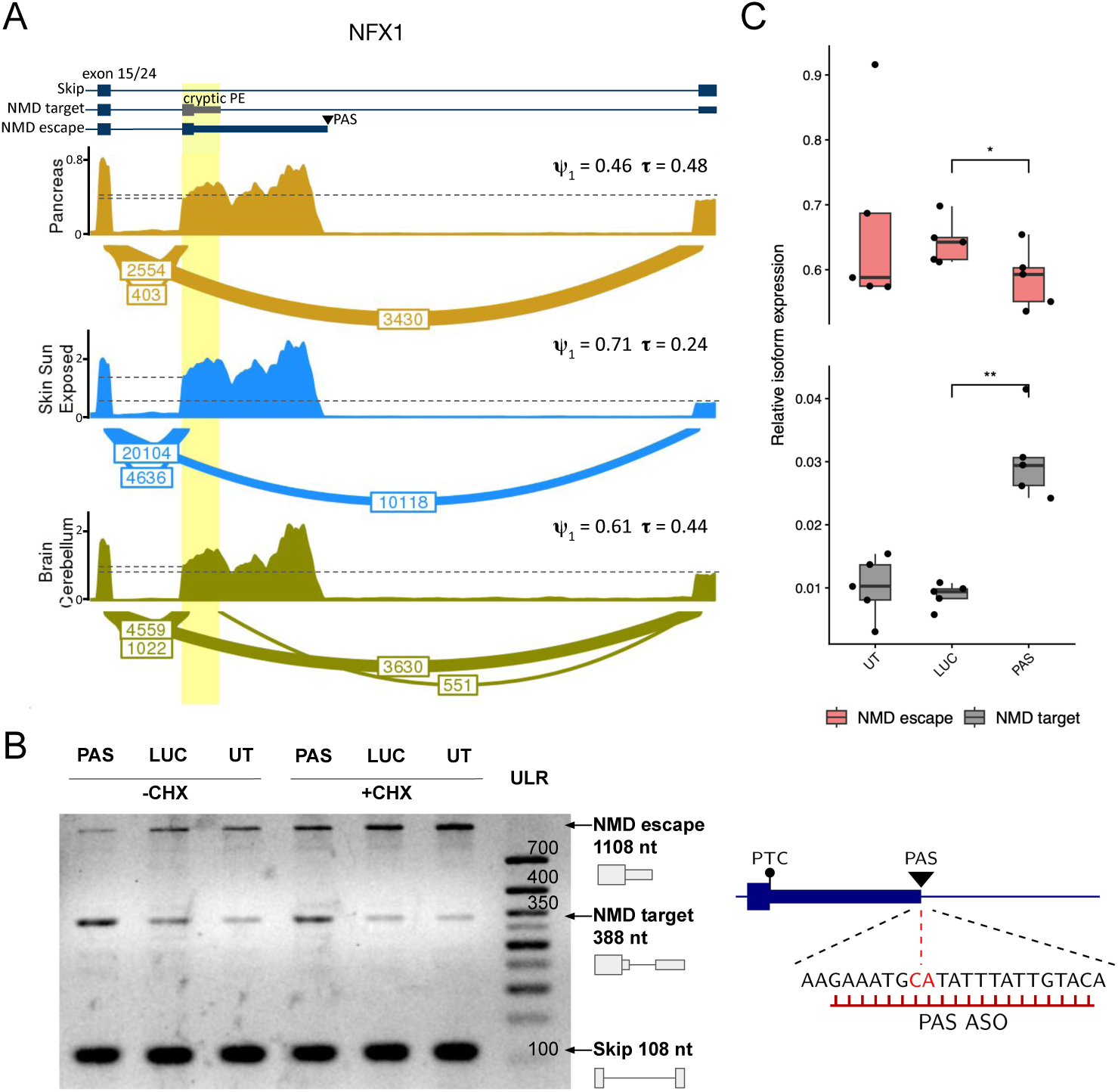
NMD escape in the *NFX1* gene. **(A)** Read coverage profiles and sashimi plots illustrating a tissue- specific switch from the skip to the NMD-escape isoform in the *NFX1* gene. The poison exon followed by a PAS is highlighted. The vertical axes and numbers on the sashimi plots are as described in Figure 3. Transcript models from CHESS are shown at the top in blue; the unannotated poison exon is shown in gray. Dotted lines indicate the drop in read coverage between the upstream and downstream exons. **(B)** Left: Endpoint RT-PCR analysis of untreated cells (UT), cells treated with ASO targeting luciferase (LUC), and ASO targeting the cleavage site (PAS), with (+CHX) or without (-CHX) cycloheximide treatment. Right: Genomic sequence surrounding the PAS targeted by the ASO. **(C)** RT-qPCR analysis showing the fraction of NMD-escape (red) and NMD-target (grey) isoforms among all three isoforms under ASO treatment in the presence of CHX. ULR — ultra low range DNA ladder.

Since an ASO that blocked only the cleavage site was sufficient to achieve the switch from the NMD-escape to the NMD-target isoform in the previous example, we designed a similar ASO for *NFX1* (Figure 4B, right). Transfection of HeLa cells with 25 nM of the ASO targeting the cleavage site promoted the inclusion of the poison exon and suppression of the NMD-escape isoform compared to transfection with the mock ASO or the untreated control, both with and without CHX treatment (Figure 4B, left). The upregulation of the NMD-target isoform and the downregulation of the NMD-escape isoform following ASO treatment were further confirmed by RT-qPCR (Figure 4D).

Next, we examined two additional Case III targets: the Notch signaling regulator *TM2D3* and the DEAD-box RNA helicase *DDX31*. TM2D3 associates with Notch1 at a region distinct from its primary ligand-binding domain, thereby enhancing its presentation on the cell surface [46]. *DDX31* encodes a nucleolar RNA helicase involved in a wide range of fundamental cellular processes and diseases, including cancer [47, 48, 49]. According to our analysis, both genes harbor cryptic poison exons located within alternative terminal exons (Figures S12A and S13A).

As observed previously, treatment of A549 cells with 25 nM or 50 nM ASO targeting the cleavage sites in these genes promoted poison exon inclusion and suppressed the NMD-escape isoform, in contrast to treatment with a mock ASO or no treatment (Figures S12B and S13B). Notably, the effect was dose dependent: higher ASO concentrations led to increased levels of the NMD-target isoform, most notably in the presence of CHX.

Interestingly, our predictions also encompass *TPM2*. This well-characterized gene encodes β-tropomyosin, which stabilizes actin filaments and regulates muscle contraction. It has been implicated in cardiac diseases, myopathies, and atherosclerosis [50, 51]. According to our findings, *TPM2* transcript isoforms that skip the poison exon predominate in the cervix, prostate, and most other tissues, whereas the NMD-escape isoform containing an alternative terminal exon is predominant in muscle, consistent with the high expression level of *TPM2* in this tissue (Figure S14A). Notably, in the heart we observe significant upregulation of the NMD-target isoform, which may help explain the reduced expression of *TPM2* in cardiac tissue [40].

Blockade of the polyadenylation signal and cleavage site in *TPM2* using ASOs did not yield the intended outcome: suppression of the NMD-escape isoform led to skipping of the poison exon, whereas the NMD-target isoform was undetectable in any of the tested cell lines (A549, Huh7, HeLa, and PC3; data not shown). However, we identified anecdotal evolutionary evidence supporting the proposed mechanism. The murine ortholog of *TPM2* contains an unannotated polyadenylation site and an unannotated poison exon at precisely aligning positions, suggesting that these elements have been evolutionarily conserved for regulatory purposes (Figure S14). However, in the absence of a specialized cardiac cell line, we chose not to investigate this gene further, despite its significant clinical relevance.

## Discussion

Early estimates suggested that up to one-third of alternatively spliced human transcripts contain a PTC and are thus susceptible to degradation by the NMD pathway [52]. However, accumulating evidence also indicates that not all PTC-containing transcripts are degraded, and a substantial fraction can undergo NMD escape [53]. For instance, a study of human genotyping datasets found that 52.3% of highly common nonsense variants can escape NMD [54]. Another study demonstrated that variability in NMD efficiency can only partially be explained by the 50-nt rule, with up to 30% of the variance remaining unexplained [55], suggesting that NMD escape is considerably more widespread than currently appreciated [14].

For a PTC-containing transcript, the two major routes of NMD escape are translation reinitiation [56, 57, 58, 59] and stop codon readthrough [60, 61, 62]. Both mechanisms enable the ribosome to traverse the mRNA, displace all EJCs, and simultaneously produce a protein from the nonsense allele. In this work, we consider yet another mechanism that also allows protein production, in which APA removes the portion of the 3’UTR signaling the presence of a PTC, thereby converting it into a normal stop codon. To date, only three genes were suggested to undergo NMD escape via this mechanism [21, 22, 20].

Our study provides strong evidence that intronic polyadenylation is a frequent mechanism of NMD escape. Part of the statistical support arises from the simple observation that annotated transcript ends are more abundant in introns following poison exons than in introns following non-poison cassette exons. Additional support comes from two novel RNA-seq-derived metrics: *φ*, which captures the imbalance between split read coverage at the 5’- and 3’-splice junctions, and *τ*, which reflects the drop in read coverage caused by intronic polyadenylation. Together, these metrics demonstrate a tendency for poison exons to be followed by an active intronic PAS (Cases I and II), suggesting that NMD escape via intronic polyadenylation may contribute to the unexplained variability in NMD efficiency. Moreover, alternative terminal exons may harbor cryptic poison exons (Case III), which remain unannotated due to NMD-mediated degradation, further expanding the repertoire of NMD escape events.

Applying stringent filtering criteria, we analyzed tissue-specific splicing and polyadenylation and identified 50 genes exhibiting tissue-specific NMD escape via intronic polyadenylation. However, this number likely represents only the tip of the iceberg, as none of the few genes previously known to undergo this mechanism were detected. The poison exon in the *HFE* gene [21] was missed because it was not annotated, and furthermore we did not observe splicing upregulation under CHX treatment when specifically examining its 3’UTR. The remaining two cases of NMD escape, in the *CSF3R* and *Tau* genes, did not pass our tissue-specific filters because they are restricted to CD34+ immune cells and Alzheimer’s disease patients, respectively. Moreover, the active NMD system in tissues limits the observable response of poison exons to APA-mediated escape, representing a major challenge in quantifying unproductive splicing rates in unperturbed physiological conditions [63, 64].

Intronic polyadenylation generates transcripts with alternative terminal exons, which can be classified into two categories: skipped terminal exons, which may function as internal terminal exons or be excluded, and composite terminal exons, which arise from polyadenylation within a retained intron [65]. The pipeline presented in this study (Figure 1) focuses on poison exons, representing only one subtype of skipped terminal exons, whereas the diversity of NMD isoforms is considerably broader. Intronic polyadenylation occurring in the intron upstream of a poison exon may generate a composite terminal exon, providing an additional route for NMD escape. Using the same methodology, it can also be shown that introns flanking poison exons at the 5’ end tend to contain more active PAS than introns flanking non-poison cassette exons. However, exploring this or other APA scenarios falls beyond the scope of the present study.

Several studies have compared the efficacy of blocking the cleavage site versus the polyadenylation signal. While both approaches are generally effective, blocking the polyadenylation signal is often slightly more efficient [66, 67]. In contrast, for *VRK3*, masking the cleavage site resulted in stronger suppression. The effectiveness of cis-element blocking can be context-dependent, influenced by partial compensation from surrounding non-canonical signals, local pre-mRNA conformation, and ASO accessibility [68]. Our results demonstrate that, for the four genes tested, blocking the cleavage site can as well be effective.

In conclusion, this study expands current knowledge of NMD escape via intronic polyadenylation from three experimentally validated cases (*HFE*, *CSF3R*, and *Tau*) to seven, with the addition of *VRK3*, *NFX1*, *TM2D3*, and *DDX31*. Further application and refinement of the analytical framework presented here should enable the identification of many additional events, thereby contributing to a more comprehensive understanding of polyadenylation-mediated NMD escape and its role in gene expression regulation.

## Authors’ contributions

DP and MV designed the study; MV performed data analysis; AK and DS performed the experiments; MV and DP wrote the first draft of the manuscript. All authors edited the final version of the manuscript.

## Acknowledgments

The authors cordially thank Prof. Vera Rybko for introducing us to this research area.

