## Supplemental material for "Novel examples of NMD escape through alternative intronic polyadenylation"

March 5, 2026

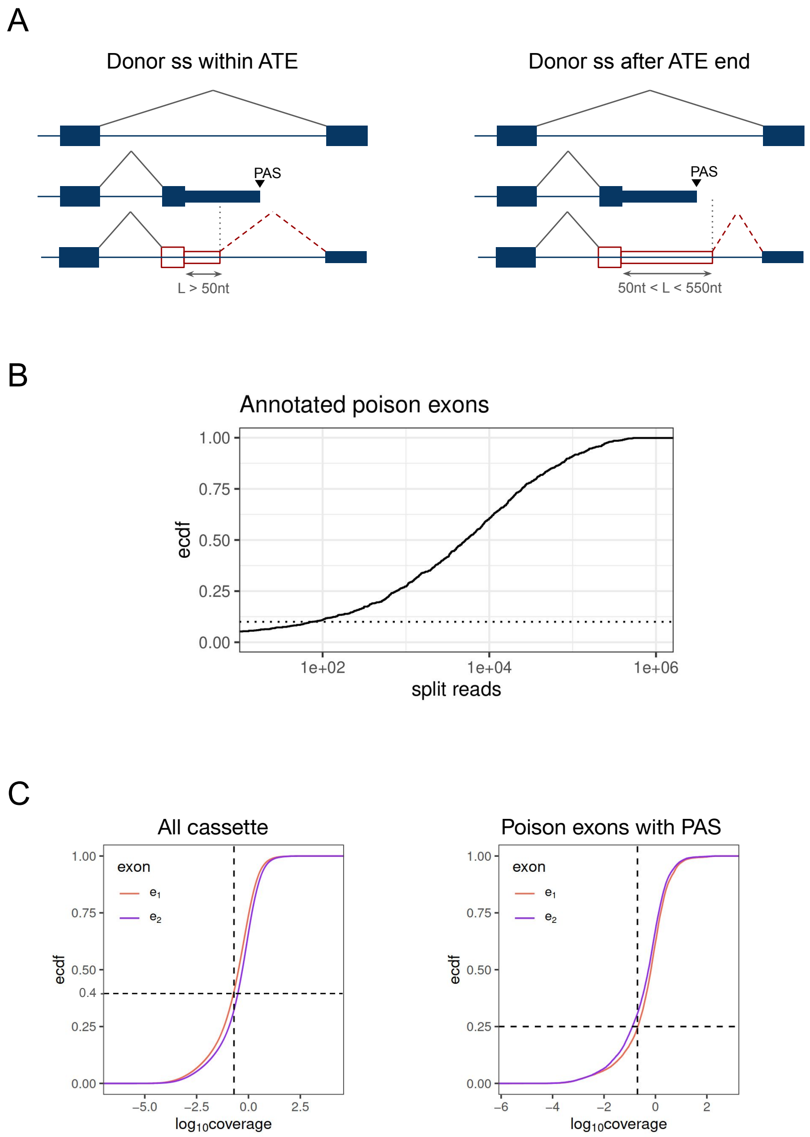

**Figure S1:** **(A)** Cryptic poison exons overlapping alternative terminal exons. Classification of cryptic poison exons according to the location of the donor splice site relative to annotated PAS. Only poison exons with fewer than 550 nucleotides between the PTC and the downstream donor splice site were considered (see Methods). Red dashed lines indicate the $I_2$ splice junction. Unannotated exons are outlined in red. **(B)** Empirical cumulative distribution function (eCDF) of split read support for $I_2$ splice junctions of annotated poison exons in GTEx. The line represents the 0.1 percentile for the annotated splice junctions, which corresponds approximately to 100 supporting reads. **(C)** eCDF of $e_1$ and $e_2$ coverage for all cassette exons (left) and poison exons with PAS (right) across GTEx tissues. The median $e_1$ value is lower than the median $e_2$ value in cassette exons and higher than the median $e_2$ value in poison exons with PAS. The $e_1 > 0.1$ threshold for $\tau$ computations excludes a quarter of exon-tissue pairs for poison exons with PAS (see Methods).

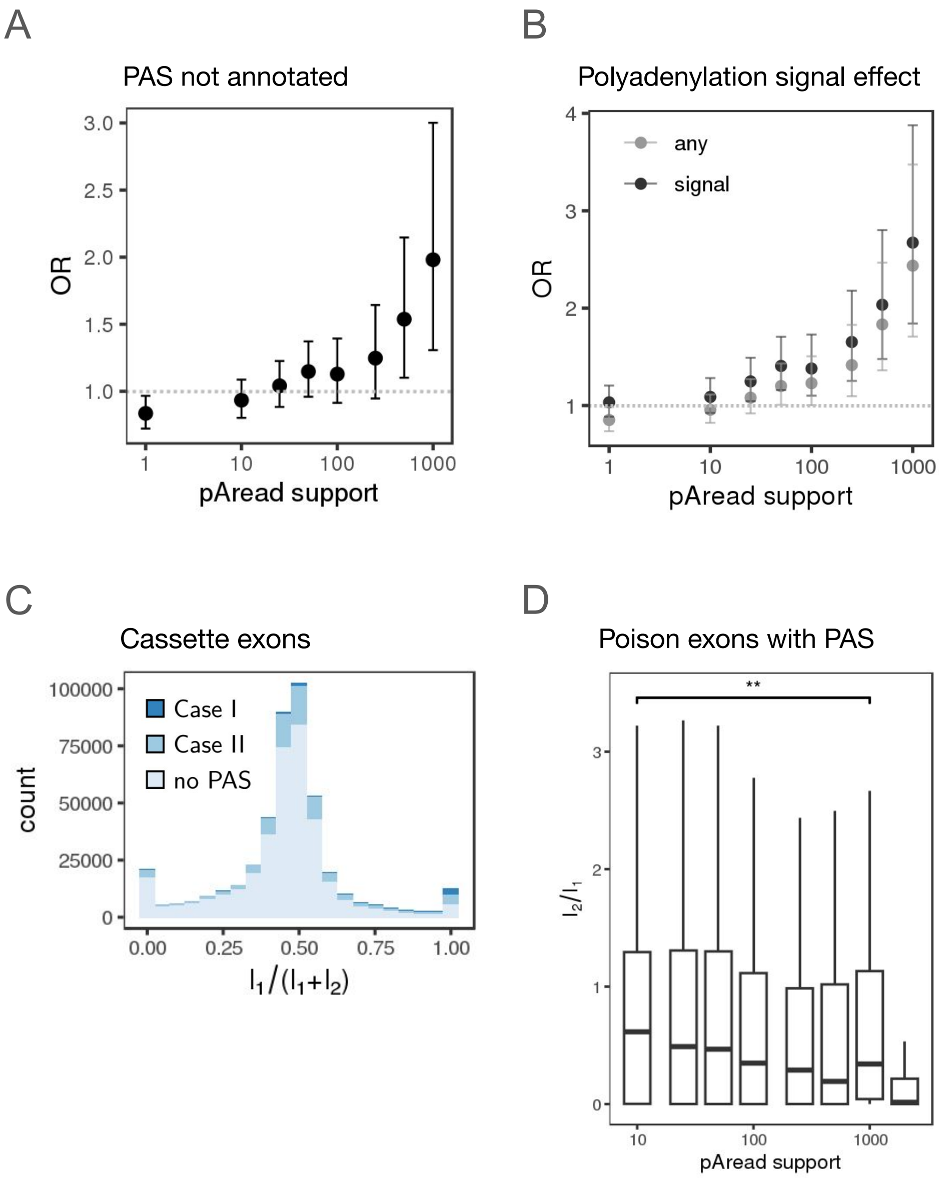

**Figure S2:** The odds ratio ($OR$) for the proportion of exons followed by a PAS in poison versus cassette exons. **(A)** $OR$ for the unannotated PAS from the polyA read-based set as a function of polyA read support. **(B)** $OR$ as a function of polyA read support for all PAS and PAS preceded by a polyadenylation signal. Whiskers represent 95% confidence intervals. **(C)** Distribution of $\varphi = I_1/(I_1 + I_2)$ values across GTEx tissues for cassette exons. Colors indicate whether the exon is followed by a PAS, as shown in Figure 1B. **(D)** Ratio of $I_2$ and $I_1$ as a function of polyA read support computed for poison exons. Asterisks denote statistical significance based on a two-sided Mann-Whitney U-test.

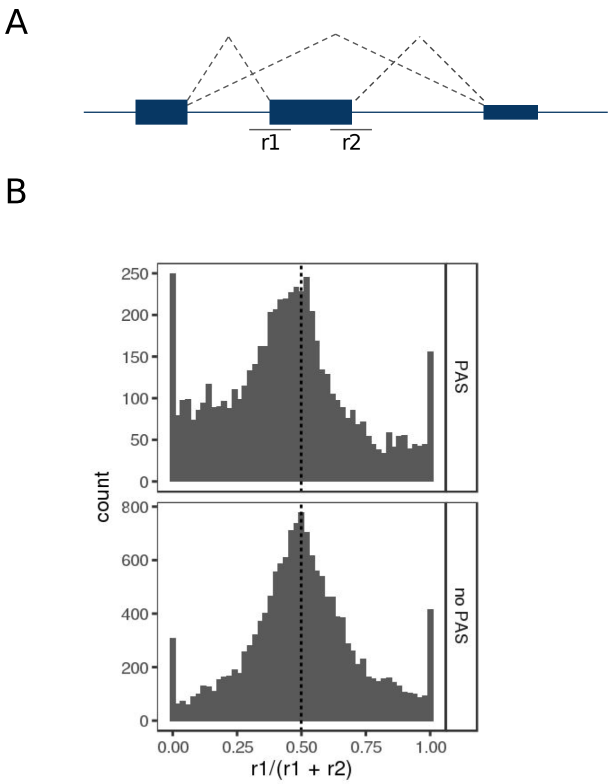

**Figure S3:** **(A)** A schematic illustrating continuous reads overlapping splice sites: $r_1$ denotes reads that overlap the 5' end of the cassette exon, and $r_2$ denotes reads that overlap the 3' end. **(B)** Distribution of the fraction of $r_1$ in the sum of $r_1$ and $r_2$ for poison exons followed by a PAS and poison exons not followed by a PAS. A low $r_1/(r_1 + r_2)$ ratio implies higher retention at the 3' end, which supports downstream PAS usage.

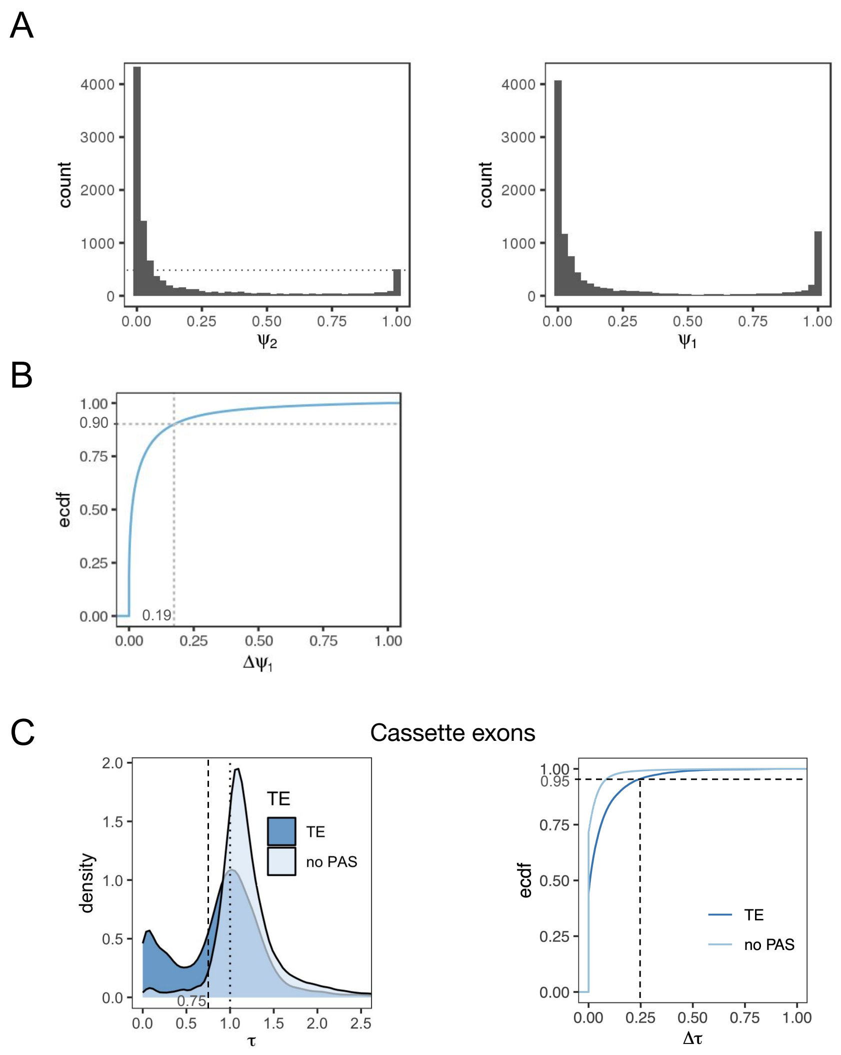

**Figure S4:** Distributions of $\psi_1$, $\psi_2$, and $\tau$ values. **(A)** Distribution of $\psi_1 = \frac{I_1}{I_1+E}$ and $\psi_2 = \frac{I_2}{I_2+E}$ for poison exons followed by a PAS in GTEx tissues. **(B)** Empirical cumulative distribution function (eCDF) of differences between $\psi_1$ values across different tissues for non-poison cassette exons not followed by a PAS. The line indicates the 90th percentile ($\Delta\psi_1 \approx 0.2$). This distribution was used to assess $\psi_1$ variability among tissues for standard tissue-specific cassette exons. The 90th percentile was set as a threshold to identify poison exons with tissue-specific inclusion of the junction upstream of the exon (Figure 2B). **(C)** left: Distribution of $\tau$ values across GTEx tissues for cassette exons followed by an annotated PAS (i.e., annotated alternative terminal exons, dark blue) and cassette exons not followed by any PAS (light blue). Exons with a PAS are clearly characterized by an additional peak at $\tau = 0$, supporting the validity of the metric. Right: eCDF of differences between $\tau$ values in different tissues for alternative terminal exons (dark blue) and cassette exons not followed by a PAS (light blue). The line shows the 95% percentile for the former set ($\Delta\tau \approx 0.25$).

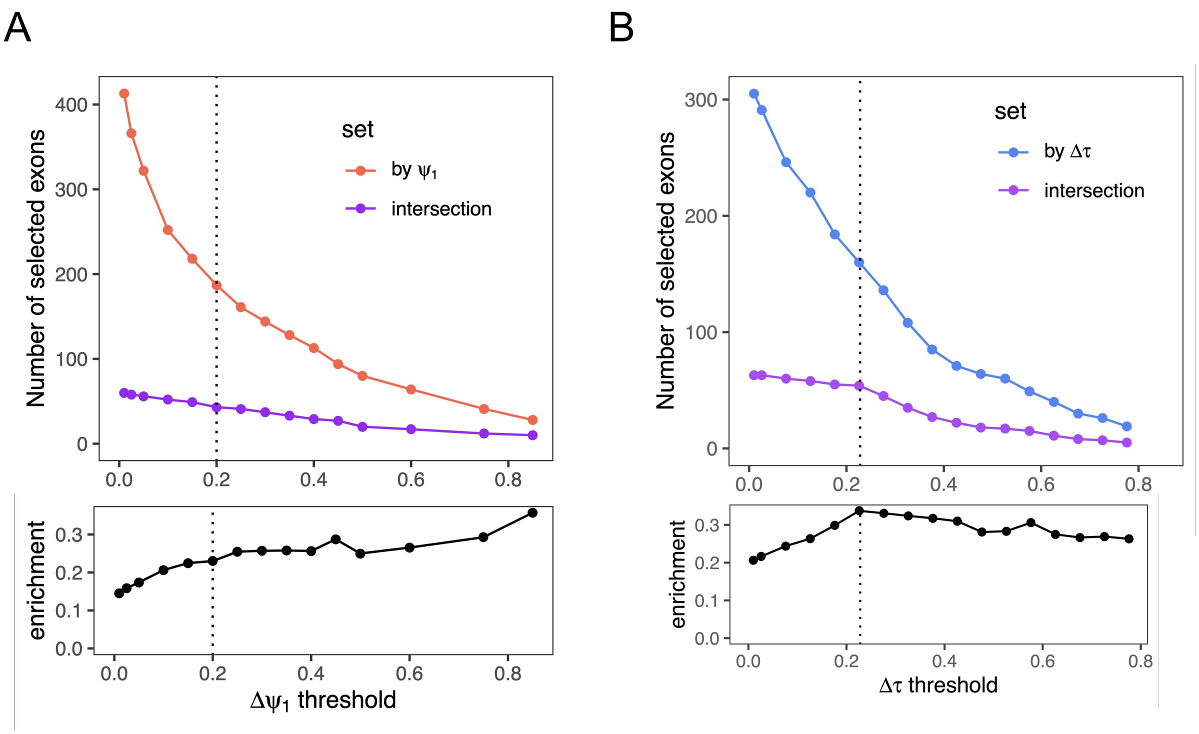

**Figure S5:** **(A)** Dependence on the splicing threshold $\Delta\psi_1$, with the polyadenylation threshold fixed at $\Delta\tau = 0.25$. The orange curve ("by $\psi_1$") represents the number of exons with $\Delta\psi_1$ above the indicated cutoff. The purple curve ("intersection") shows the subset that additionally satisfies $\Delta\tau \geq 0.25$ and exhibits a negative cross-tissue correlation between $\psi_1$ and $\tau$. The bottom panel reports enrichment, defined as(intersection)/(by $\psi_1$). **(B)** Dependence on the polyadenylation threshold $\Delta\tau$, with the splicing threshold fixed at $\Delta\psi_1 = 0.2$. The blue curve ("by $\Delta\tau$") represents the number of exons with $\Delta\tau$ above the indicated cutoff. The purple curve ("intersection") shows the subset that additionally satisfies $\Delta\psi_1 \geq 0.2$ and exhibits a negative correlation between $\psi_1$ and $\tau$. The bottom panel reports enrichment, defined as (intersection)/(by $\Delta\tau$). Dotted vertical lines indicate the thresholds used in the analysis: $\Delta\psi_1 = 0.2$ and $\Delta\tau = 0.25$.

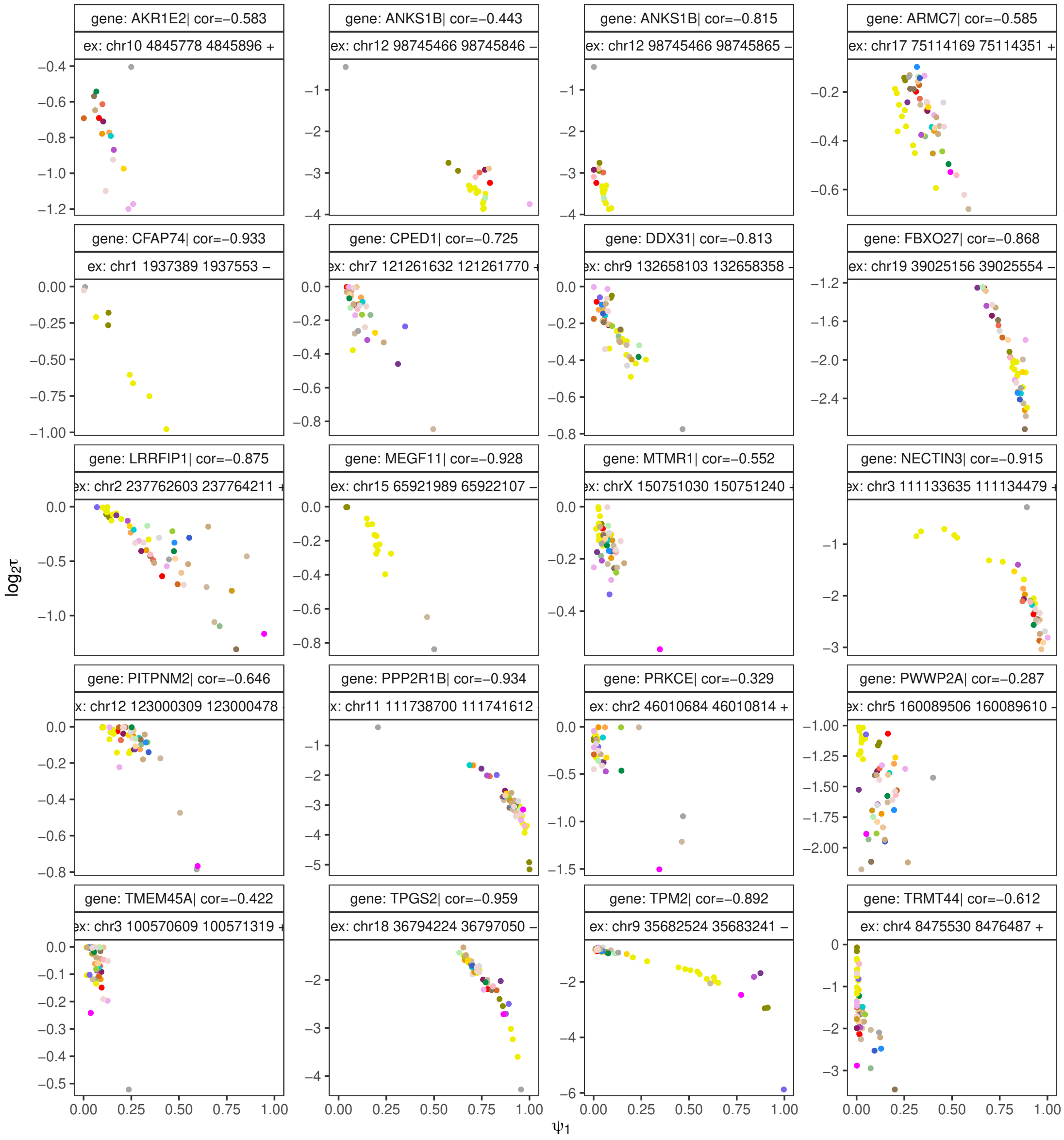

**Figure S6: Part 1 — continued on the next page.**

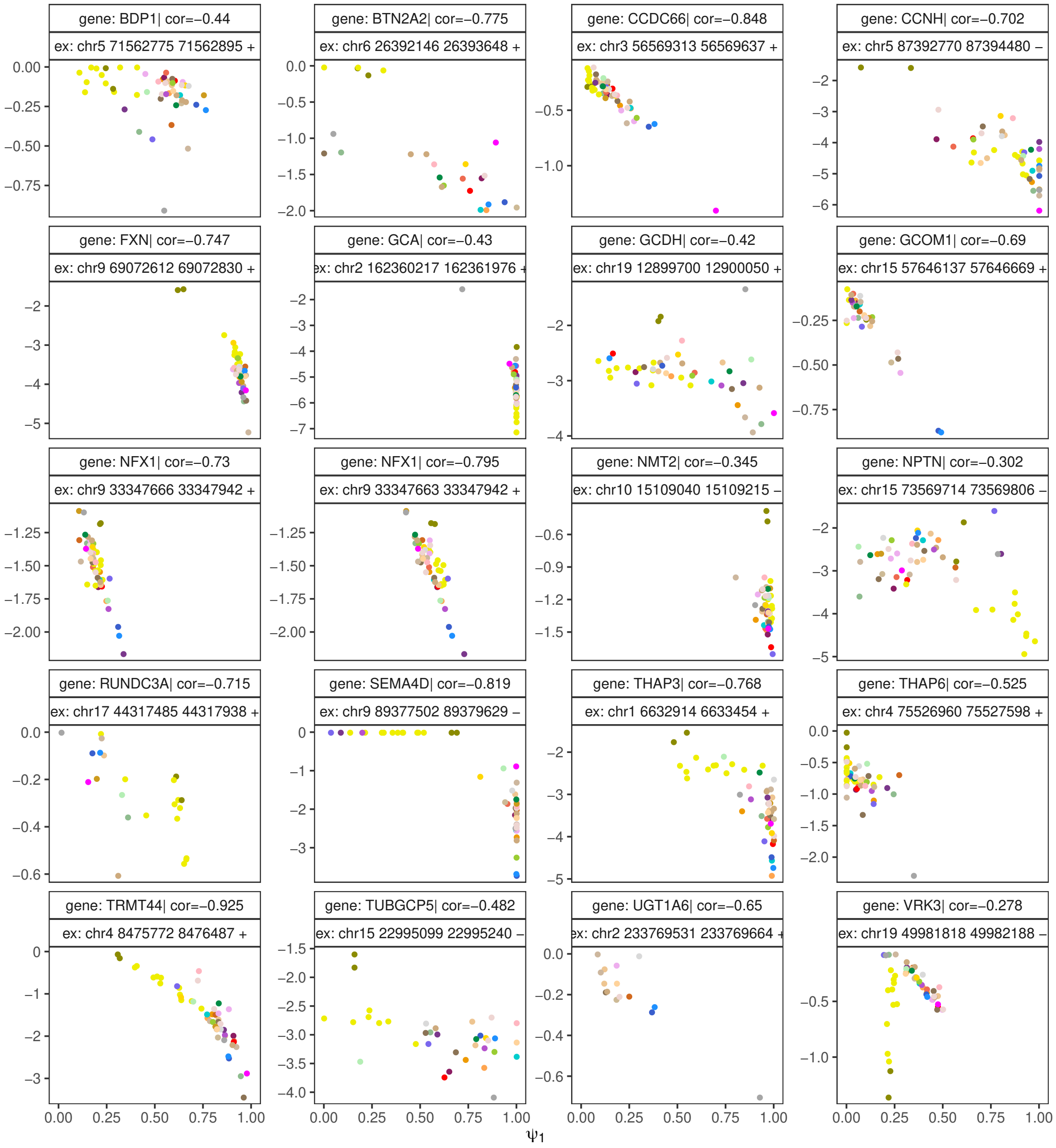

**Figure S6: Part 2 — continued on the next page.**

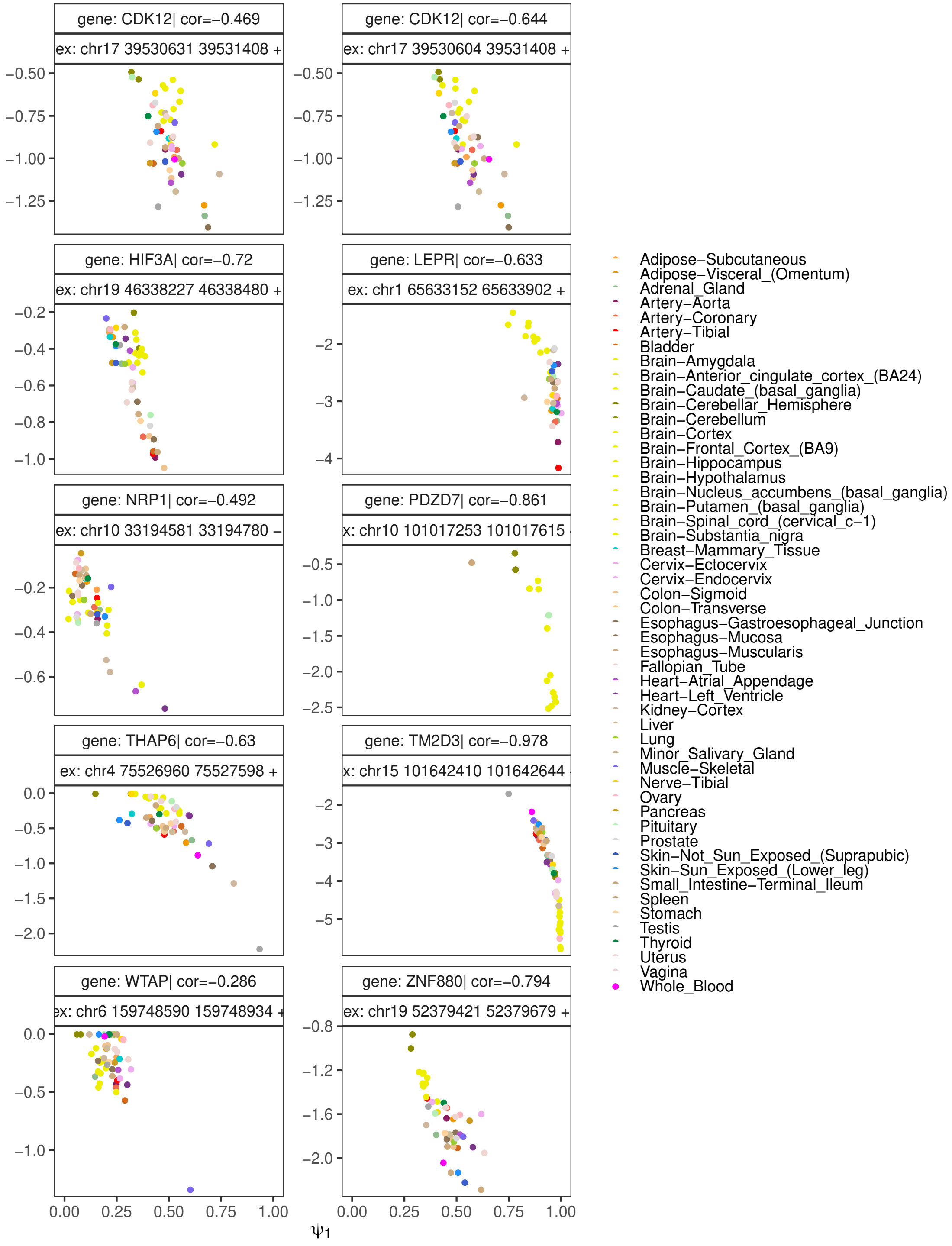

**Figure S6: Part 3 — continued from the previous page.** Mean $\tau$ and $\psi_1$ values across GTEx tissues for the selected poison exons exhibiting a tissue-specific switch between skip and NMD-escape isoforms. Poison exon coordinates, gene name, and the Spearman correlation coefficient between $\psi_1$ and $\tau$ are displayed above each dot plot.

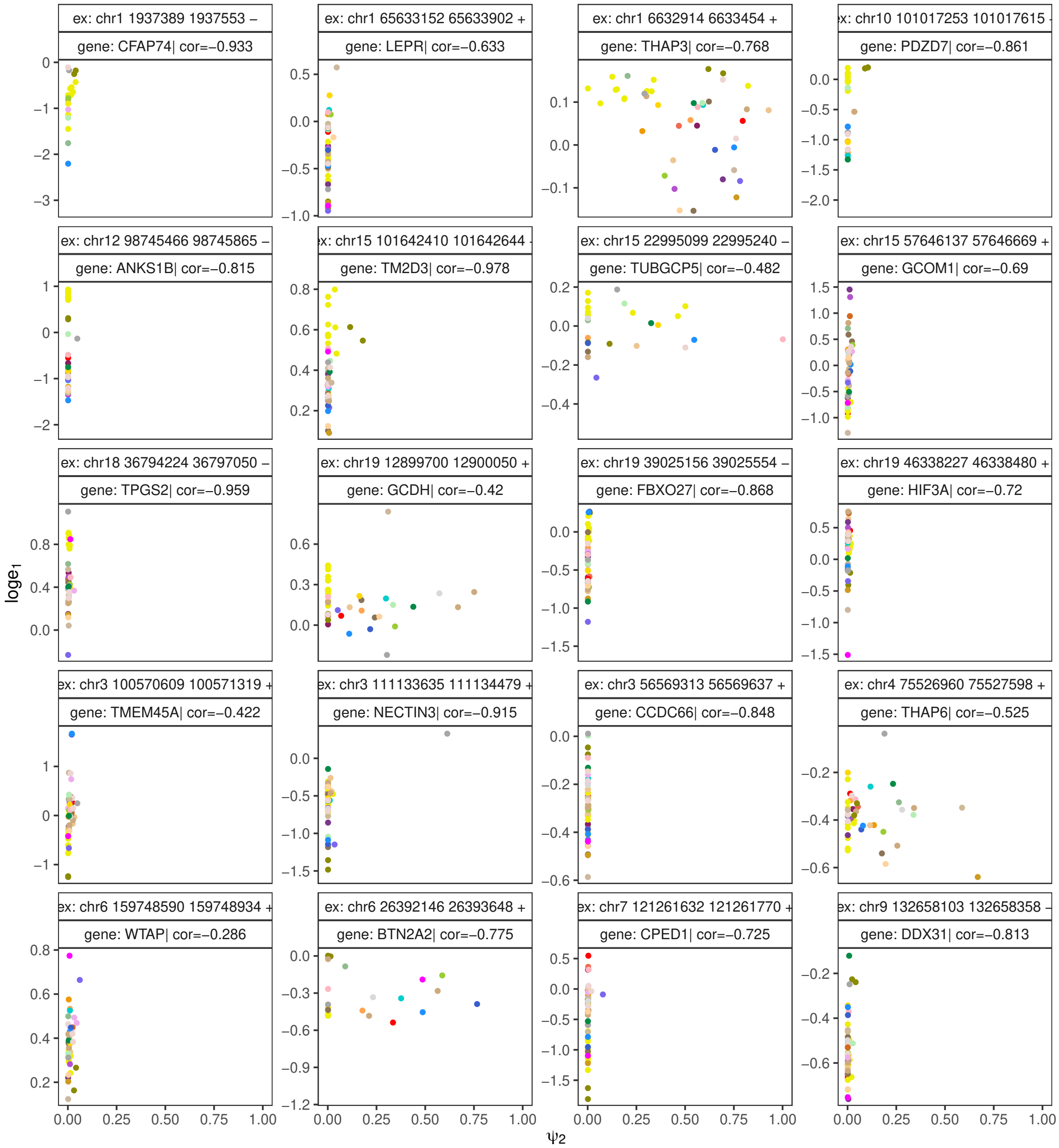

**Figure S7: Part 1 — continued on the next page.**

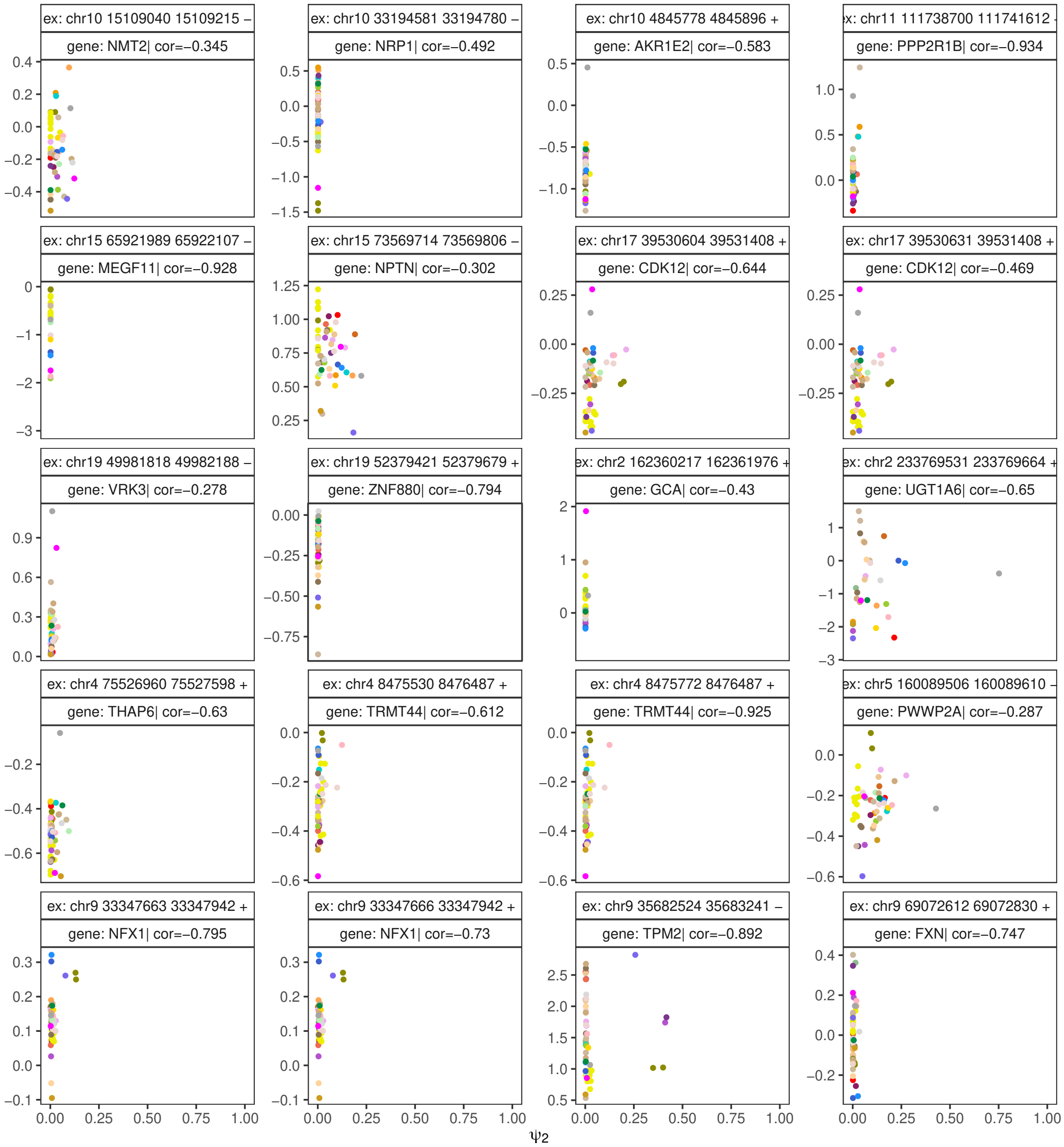

**Figure S7: Part 2 — continued on the next page.**

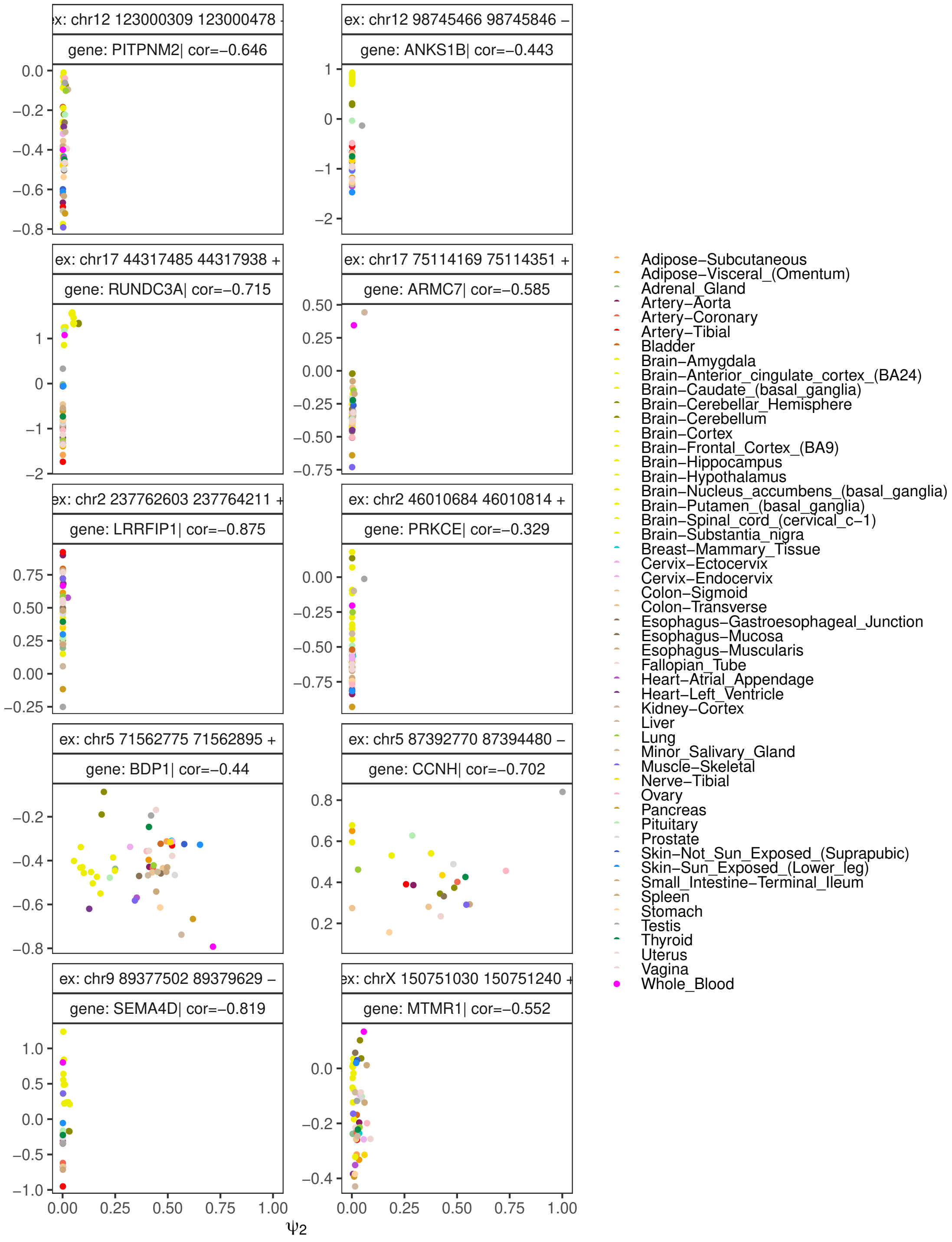

**Figure S7: Part 3 — continued from the previous page.** Mean $\tau$ and upstream exon coverage ($e_1$) across GTEx tissues for the selected poison exons exhibiting a tissue-specific switch between skip and NMD-escape isoforms. Poison exon coordinates, gene name, and the Spearman correlation between gene expression in TPM and $\tau$ are displayed above each dot plot.

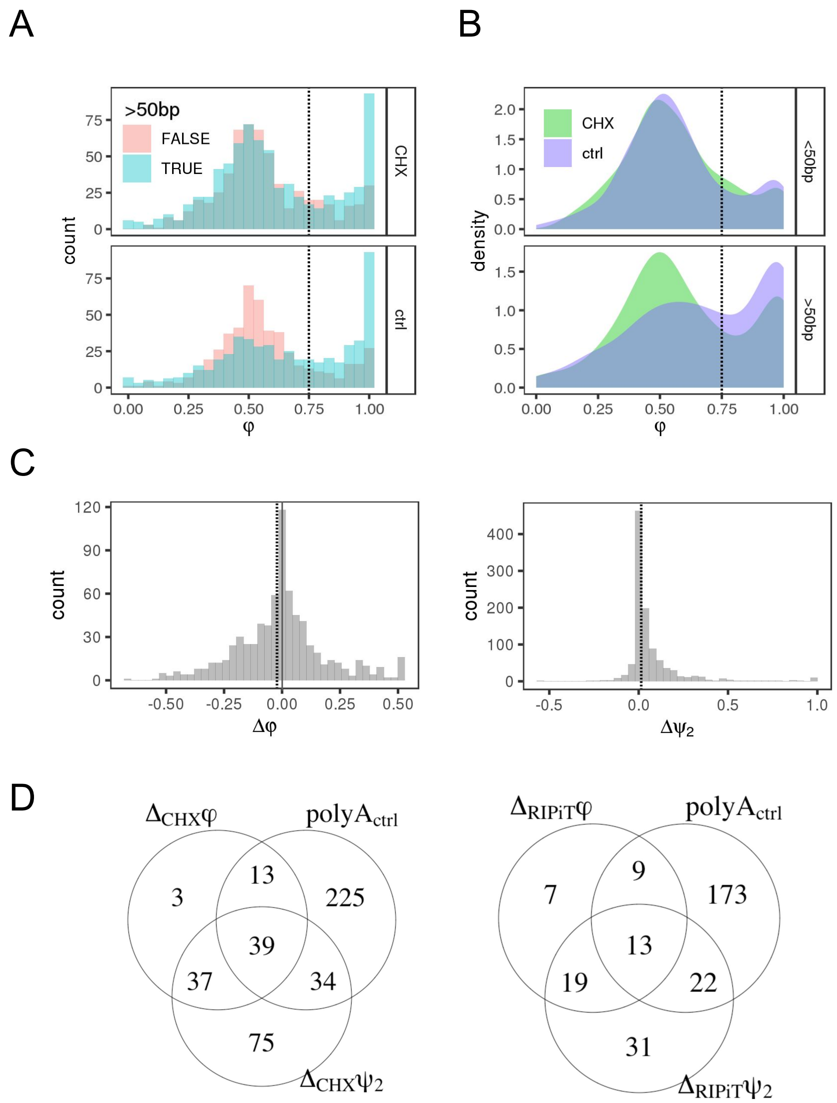

**Figure S8:** Response of cassette exons followed by a PAS to NMD inactivation. **(A,B)** Distribution of $\varphi$ values for exons with a stop codon followed by a PAS before and after NMD inhibition with CHX. In (A), colors indicate whether the exon obeys the 50-nt rule (191 of 361 exons do). In (B), colors indicate whether the cells were treated with CHX. **(C)** Distribution of changes in $\varphi$ and $\psi_2$ upon NMD inhibition: $\Delta\psi_2 = \psi_2^{CHX} - \psi_2^{ctrl}$, $\Delta\varphi = \varphi^{CHX} - \varphi^{ctrl}$, for poison exons followed by a PAS. Dashed lines indicate thresholds used to select candidates expressing both NMD-escape and NMD-target isoforms $\Delta\varphi \leq -0.025$, $\Delta\psi_2 \geq 0.015$ and to indicate polyadenylation in untreated cells $\varphi_{ctrl} \geq 0.75$ or $\delta_{ctrl} \leq 0.7$. **(D)** Venn diagrams illustrating the response of poison exons followed by a PAS to NMD inactivation. The category $polyA_{ctrl}$ denotes exons with high $\varphi_{ctrl}$ or low $\tau_{ctrl}$ values in untreated cells. $\Delta$ represents the changes in the respective metrics between NMD inactivation and control conditions (see Methods). The left diagram corresponds to the dataset compiled from four cell lines treated with CHX, whereas the right diagram is based on the RIPiT dataset (see Methods).

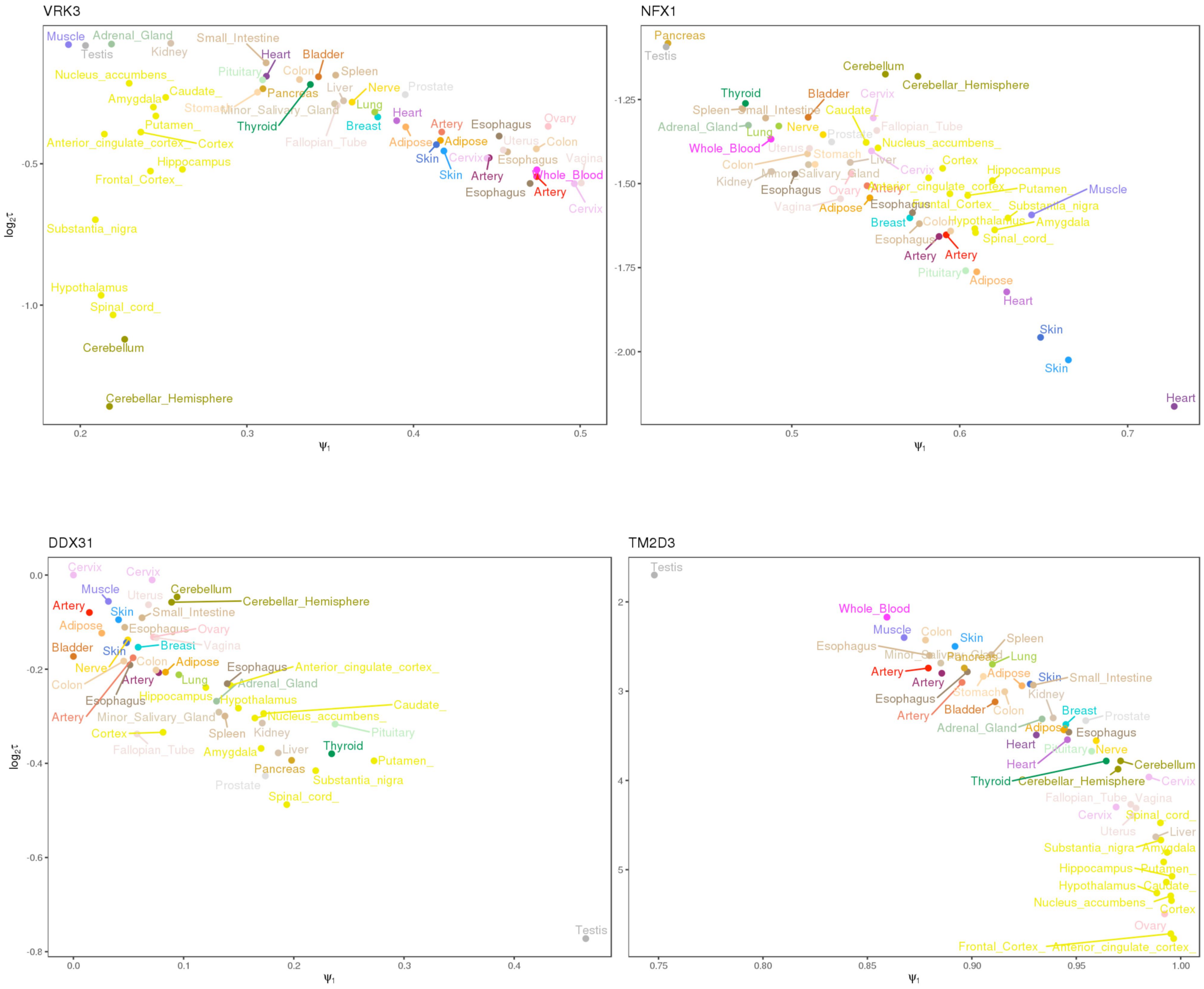

**Figure S9:** Correlations between $\tau$ and $\psi_1$ for *VRK3*, *NFX1*, *TM2D3*, and *DDX31* genes. Yellow and dark green colors correspond to different brain regions.

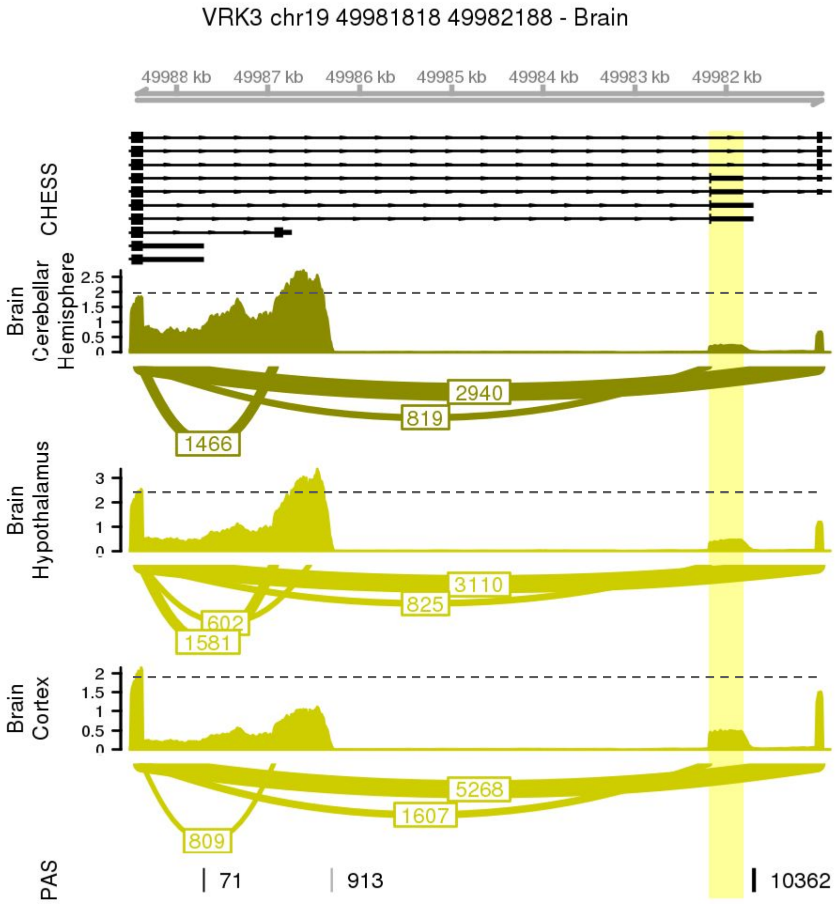

**Figure S10:** Read coverage profiles and sashimi plots for three brain tissues showing the poison exon in the *VRK3* gene. As in Figure 3, the poison exon followed by a PAS is highlighted. The vertical axes represent read coverage density, and the numbers on the sashimi plots indicate split read counts. Transcript models from CHESS are shown at the top. The PAS track displays the number of polyA reads supporting each PAS. Dotted lines illustrate the drop in read coverage between the upstream and downstream exons.

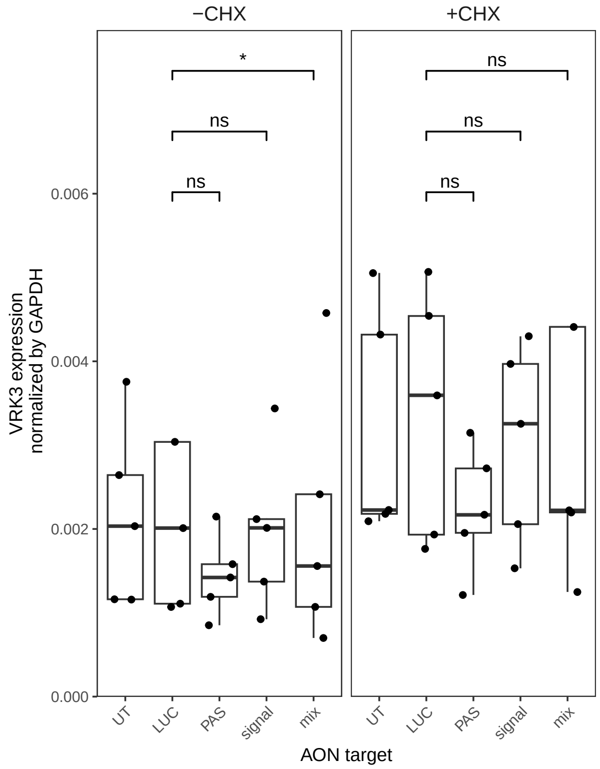

**Figure S11:** Total *VRK3* expression under ASO treatment, with and without NMD inhibition. Isoform-specific qRT-PCR results were normalized to GAPDH expression. An asterisk (*) indicates a statistically significant difference at the 5% level; “ns” denotes no statistically significant difference (one-sided Wilcoxon signed-rank test, $n = 5$ biological replicates).

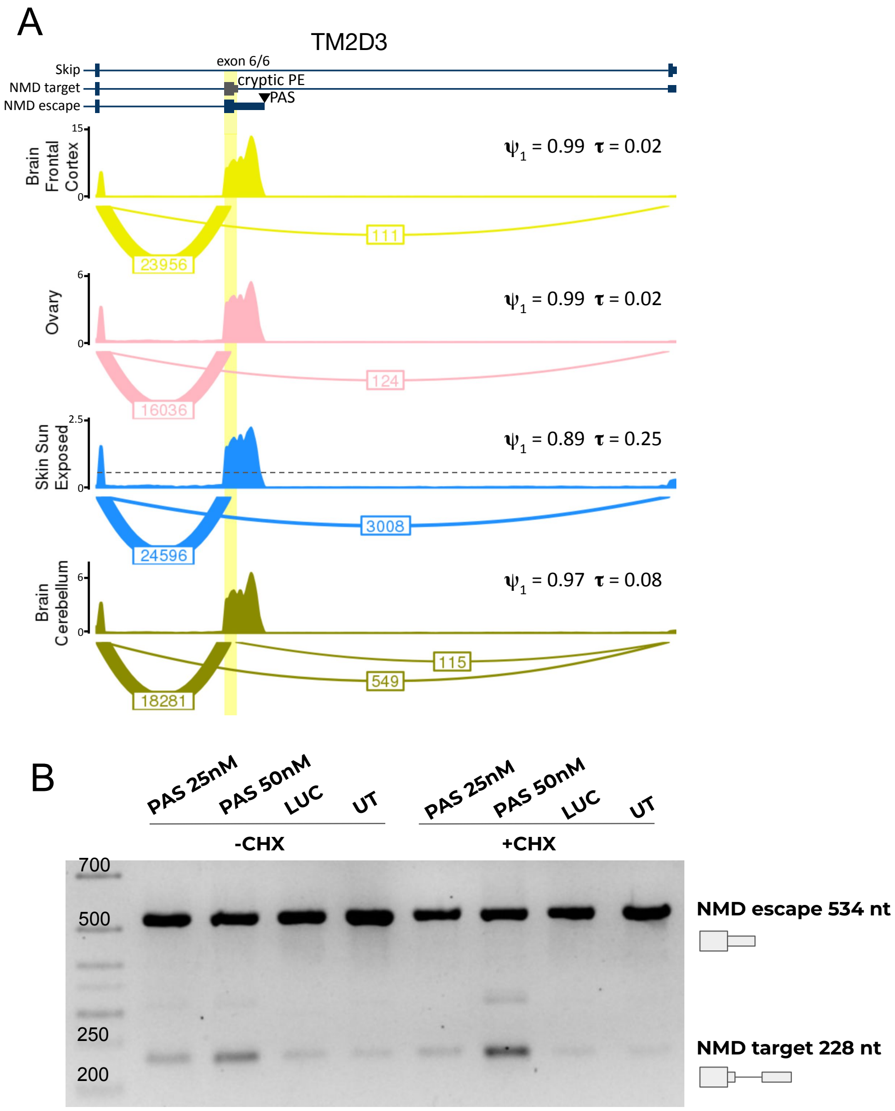

**Figure S12:** NMD escape in the *TM2D3* gene. **(A)** Read coverage profiles and sashimi plots illustrating a tissue-specific switch from the NMD-escape to the skip isoform in the *TM2D3* gene. The poison exon followed by a PAS is highlighted. The vertical axes represent read coverage density, and the numbers on the sashimi plots indicate split read counts summed across samples from each tissue. Transcript models from CHESS are shown at the top in blue, with the unannotated poison exon shown in gray. Dotted lines indicate the drop in read coverage between the upstream and downstream exons. **(B)** Endpoint RT-PCR analysis of untreated cells (UT), cells treated with ASO targeting luciferase (LUC), and ASO targeting the cleavage site (PAS) at two concentrations (25 and 50 nM), with (+CHX) or without (-CHX) cycloheximide treatment.

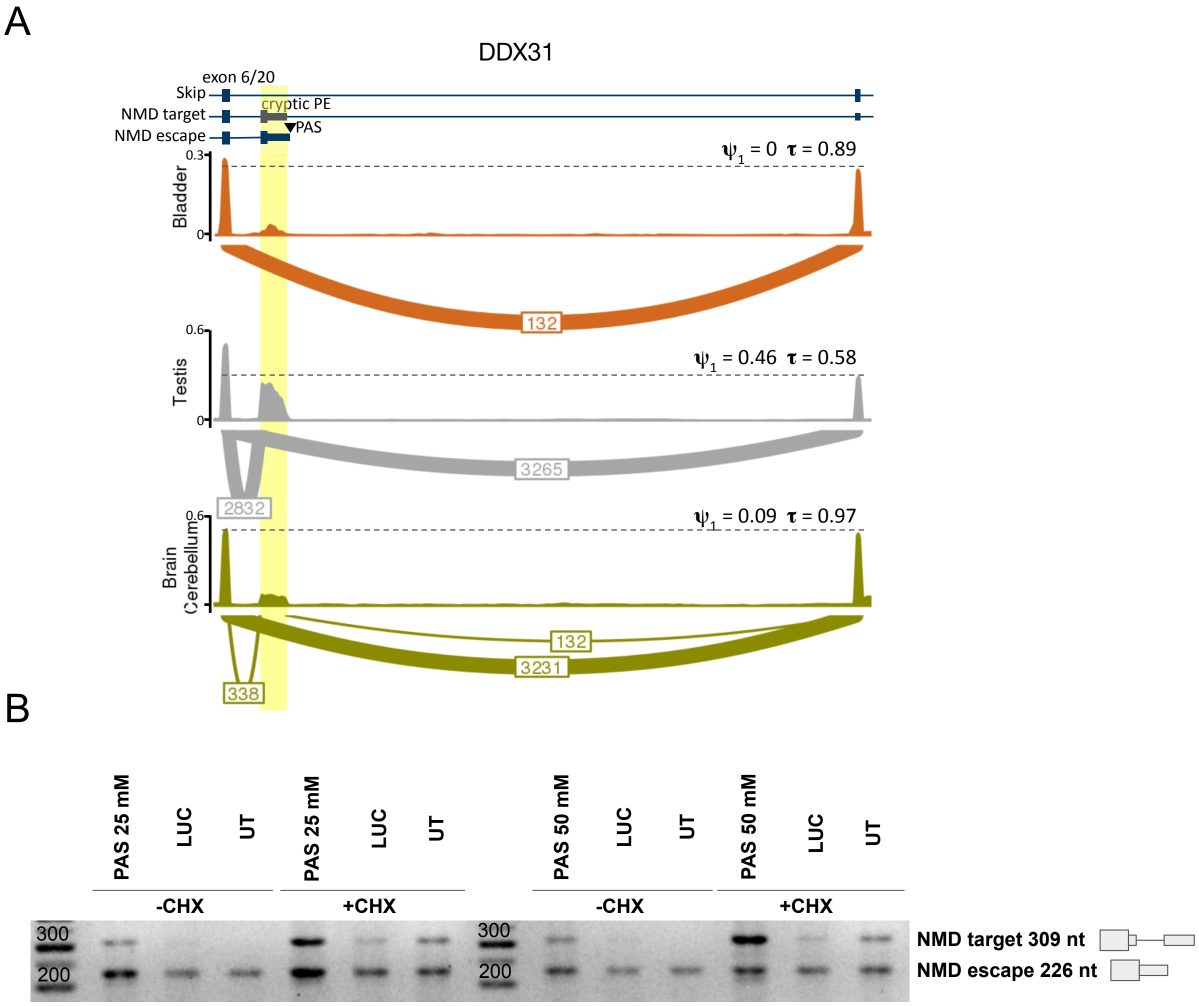

**Figure S13:** NMD escape in the *DDX31* gene. **(A)** Read coverage profiles and sashimi plots illustrating a tissue-specific switch from NMD-escape to skip isoform in the *DDX31* gene. The poison exon followed by the PAS is highlighted. Vertical axes, numbers on the sashimi plots, and transcript models are as described in Figure S12. Dotted lines indicate the drop in read coverage between the upstream and downstream exons. **(B)** Endpoint RT-PCR for untreated cells (UT) and cells treated with ASOs against luciferase (LUC) or the cleavage site (PAS) at two concentrations (25 and 50 nM), with (+CHX) and without (-CHX) cycloheximide treatment.

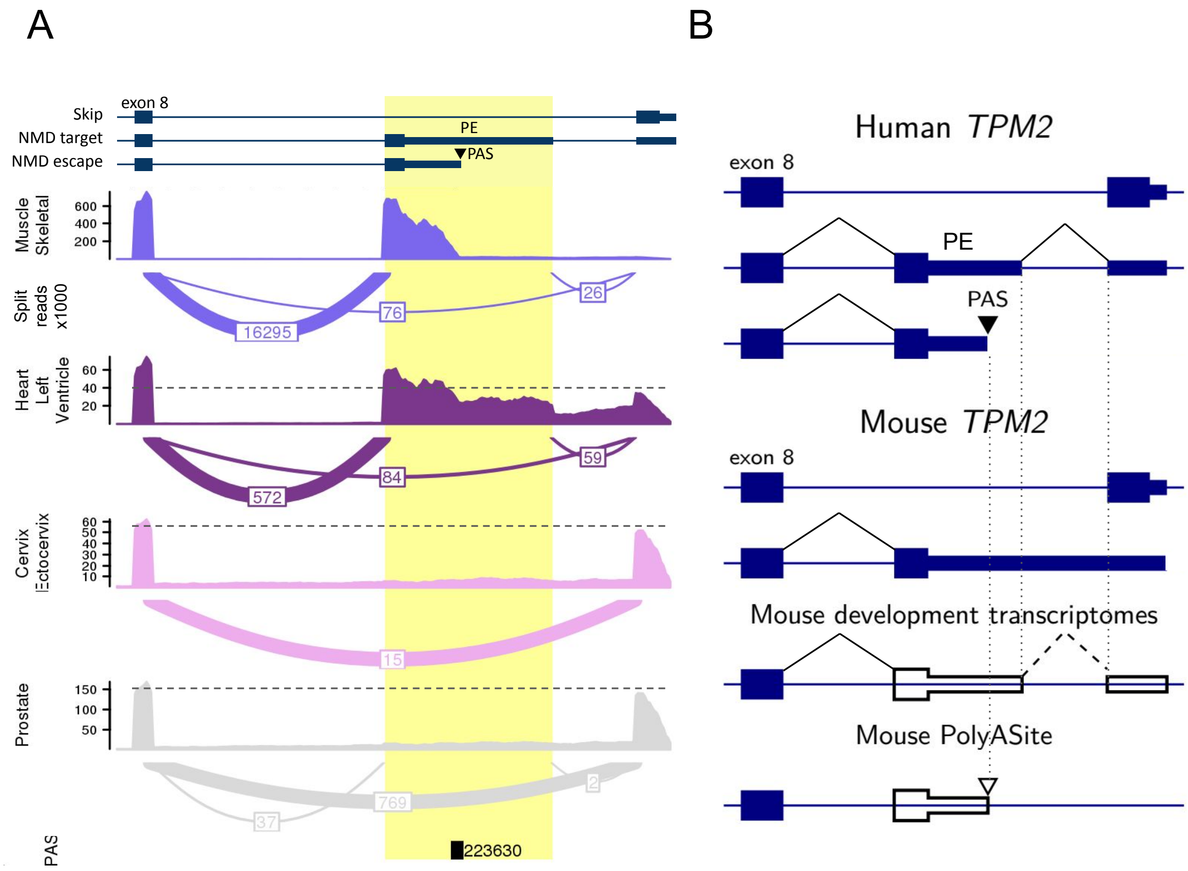

**Figure S14:** **(A)** Read coverage profiles and sashimi plots illustrating the predicted tissue-specific switch from NMD-escape to skip isoform in the *TPM2* gene. The poison exon followed by a PAS is highlighted. The vertical axes, numbers on the sashimi plots, and transcript models are as described in Figure S12. The PAS track indicates the number of poly(A) reads supporting each PAS. **(B)** Top: schematic representation of CHESS transcript models for the human *TPM2* gene. Middle: GENCODE (vM25) transcript models for the mouse *TPM2* gene. Bottom: transcript models derived from split reads in mouse embryonic development transcriptomes (ENCODE portal ID: ENCSR574CRQ) and from PAS in the mouse PolyASite database. The poison exon followed by the PAS is indicated (PE). Vertical dotted lines denote homologous positions. Unannotated exons are outlined, and dashed lines indicate unannotated splice junctions.

| ASO | Sequence |
|---|---|
| *VRK3* signal | mU*mA*mA*mA*mG*mA*mU*mU*mU*mU*mA*mU*mU*mA*mC*mU*mU*mA*mC*mA* |
| *VRK3* PAS | mG*mA*mG*mG*mA*mA*mA*mG*mA*mU*mC*mC*mA*mA*mG*mA*mU*mG*mG*mA* |
| *NFX1* PAS | mG*mU*mA*mC*mA*mA*mU*mA*mA*mA*mU*mA*mU*mG*mC*mA*mU*mU*mU*mC* |
| *TM2D3* PAS | mG*mC*mC*mC*mA*mC*mA*mG*mU*mA*mU*mU*mU*mC*mU*mU*mA*mA*mC*mU* |
| *DDX31* PAS | mA*mG*mC*mA*mU*mG*mA*mA*mU*mG*mG*mA*mU*mA*mG*mC*mU*mG*mC*mA* |
| LUC 2'OMe | mC*mU*mU*mA*mC*mG*mC*mU*mG*mA*mG*mU*mA*mC*mU*mU*mC*mG*mA* |

**Table S1:** ASO sequences. RNA bases: G, A, U, and C. 2'-OMe bases: mG, mA, mU, and mC. Phosphorothioated RNA bases: G*, A*, U*, and C*.

| Isoform | Primer | Sequence |
|---|---|---|
| Skip isoform | VRK3_F | AACAGAAGTTTGTTGATAAGCCGG |
| | VRK3_R | CTGAGGGCCTGATCCAGT |
| | NFX1_F | CAGAAGTGGTGCATGGGCAA |
| | NFX1_R | GTGCATCCCACATGGTAGCG |
| NMD-target isoform | VRK3_F_nmd | TCCCTGACCAGTGTTTGTTGAT |
| | VRK3_R_nmd | CTGAGGGCCTGATCCAGT |
| | NFX1_F_nmd | ATGGAAAACCAAACATCTTTCGGAG |
| | NFX1_R | GTGCATCCCACATGGTAGCG |
| NMD-escape isoform | VRK3_F_polyA | CACGAGCTCCCTGACCA |
| | VRK3_R_polyA | TGACCTCCCCACCCGT |
| | NFX1_F_polyA | TGTAAAAAATGTGATATCTATGAAATGC |
| | NFX1_R_polyA | TTTTTTTTTTTTTGCATTTCTTATTGGACTTC |

**Table S2:** Primers for qPCR.

| Primer | Sequence |
|---|---|
| VRK3_F_polyA | CACGAGCTCCCTGACCA |
| VRK3_R_nmd | TGTTCCTCAGCATGGCGTAG |
| VRK3_R_polyA | TGACCTCCCCACCCGT |
| NFX1_F | CAGAAGTGGTGCATGGGCAA |
| NFX1_R | GTGCATCCCACATGGTAGCG |
| NFX1_R_polyA | TTTTTTTTTTTTTGCATTTCTTATTGGACTTC |
| TM2D3_F_PE | ACGTCCTGCTCATTGGAGTT |
| TM2D3_R | GGACGCTCATCTGCAACAAC |
| TM2D3_R_polyA | CACATGACTGAAGCTGAGCC |
| DDX31_F_polyA | TCCTGGCTCCTGTGCTATGT |
| DDX31_F_PE | GCCATGCAGCAAGTAATGGAG |
| DDX31_R | ACACTCCAGGCACAATCCAG |
| DDX31_R_polyA | TTTTTTTTTTTTTAGCTGCAAAACCAAATCAC |

**Table S3:** Primers for endpoint PCR (skip, NMD-target and NMD-escape isoforms).
